# Laser Ablation and Fluid Flows Show a Single Force Mechanism Governs Spindle Positioning

**DOI:** 10.1101/2021.11.21.469320

**Authors:** Hai-Yin Wu, Gökberk Kabacaoğlu, Ehssan Nazockdast, Huan-Cheng Chang, Michael J. Shelley, Daniel J. Needleman

## Abstract

Few techniques are available for elucidating the nature of forces that drive subcellular behaviors. Here we develop two complementary ones: 1) femtosecond stereotactic laser ablation (FESLA), which rapidly creates complex cuts of subcellular structures, thereby allowing precise dissection of when, where, and in what direction forces are generated; and 2) assessment of subcellular fluid flows, by comparing direct flow measurements, using microinjected fluorescent nanodiamonds, to large-scale fluid-structure simulations of different models of force transduction. We apply these to study centrosomes in *Caenorhabditis elegans* early embryos, and use the data to construct a biophysically-based model of centrosome dynamics. Taken together, we demonstrate that cortical pulling forces provide a general explanation for many behaviors mediated by centrosomes, including pronuclear migration/centration and rotation, metaphase spindle positioning, asymmetric spindle elongation and spindle oscillations. In sum, this work establishes new methodologies for disentangling the forces responsible for cell biological phenomena.

## INTRODUCTION

Forces are crucial drivers of subcellular organization, determining both the movement of subcellular structures and their stability once positioned. While a range of sophisticated tools have been developed to measure the magnitude of such forces (Roca-Cusachs et al., 2017), and genetic knockouts can be used to identify their molecular bases, the mechanical origins of cell biological forces remains difficult to study. For example, molecular perturbations demonstrate that dynein is required for aster motion in diverse systems, but it remains controversial if the relevant forces are based on aster interactions with bulk cytoplasm or with the cell cortex (Wu et al., 2017, Xie and Minc, 2020).

The movement and positioning of centrosomes – microtubule (MT) organizing centers that are often at the middle of asters in cells - govern many important phenomena in cell biology. This includes: the orientation and positioning of the spindle, which controls the orientation and positioning of the division plane in many contexts (Knoblich, 2010, Kotak et al., 2012, Siller and Doe, 2009, Kiyomitsu and Cheeseman, 2012, Pearson and Bloom, 2004, Tame et al., 2014, von Dassow et al., 2009, Minc et al., 2011); the position of the nucleus, which can strongly influence interphase subcellular organization (Reinsch and Gonczy, 1998, Reinsch and Karsenti, 1997, Rujano et al., 2013); the migration of pronuclei, which unite the maternal and paternal genetic material (Longo and Anderson, 1968, Meaders and Burgess, 2020). Centrosomes display three basic behaviors in various systems: **1)** directed motion (Cowan and Hyman, 2004, von Dassow et al., 2009, Longo and Anderson, 1968, Meaders and Burgess, 2020, Tanimoto et al., 2018), **2)** stable positioning (Garzon-Coral et al., 2016, Foe and von Dassow, 2008, von Dassow et al., 2009), and **3)** oscillations (Du and Macara, 2004, Riche et al., 2013, Zhu et al., 2013). All of these behaviors are observed in early *C. elegans* embryos: the pronuclear complex (PNC) undergoes directed motion to the cell center with its two centrosomes becoming aligned with the embryo’s long axis (Cowan and Hyman, 2004). The spindle then forms between the two centrosomes and is stably positioned in the cell center (Garzon-Coral et al., 2016). Finally, as the spindle elongates and is positioned asymmetrically, it and its centrosomes undergo transverse oscillations (Cowan and Hyman, 2004, Pecreaux et al., 2006, Riche et al., 2013, Grill et al., 2005, Grill et al., 2001, Grill et al., 2003). It is poorly understood how these different behaviors arise from underlying mechanical and biochemical processes, though extensive work demonstrates the importance of astral MTs that radiate outward from the centrosomes, and of the molecular motor dynein (Colombo et al., 2003, Bringmann et al., 2007, Goulding et al., 2007, Galli et al., 2011, Tsou et al., 2002, Nguyen-Ngoc et al., 2007, Krueger et al., 2010, Gotta et al., 2003).

All movements in cells result from forces. Due to the small size of the centrosomes (<10 μm), their slow speeds (<1 μm/sec), and the viscous nature of the cytoplasm (>100 mPa·sec), the Reynolds number associated with their motion is Re < 10^−7^ << 1, thus inertial forces are negligible in comparison to viscous forces. Consequently, the velocity of a centrosome is proportional to the forces acting upon it. Thus, a central challenge is to identify the origin of those forces yielding these distinct behaviors of centrosomes (i.e. directed motions, stable positioning, and oscillations). Most proposals center around three possible types of forces acting through astral MTs: MTs pushing on the cortex, MTs pulled by force generators anchored to the cortex, or MTs pulled by force generators in the cytoplasm (Wu et al., 2017).

Here, we developed biophysical approaches to unambiguously determine the extent to which different centrosome motions in one-cell *Caenorhabditis elegans* embryos are driven by MT pushing or by pulling, and if pulling, of cortical or cytoplasmic origins. This relies on several advances. Firstly, we developed and utilized femtosecond stereotactic laser ablation (FESLA) as a versatile tool to dissect when, where, and in what direction forces originate. From our ablation studies, we conclude that pulling forces dominate during PNC migration and rotation, spindle centering, elongation, and oscillations. Secondly, at all these stages we compared subcellular flow measurements using microinjected fluorescent nanodiamonds (FNDs) with large-scale fluid dynamics simulations of flows resulting from pulling forces on astral MTs, from either force generators residing on the cortex or in the cytoplasm. The simulations of the former showed remarkable agreement with flow measurements, while those of the latter were profoundly different. This supports the hypothesis of pulling originating from the cortex, while also demonstrating that subcellular flows can encode in their structure a powerful signature of the mechanical basis of force generation. Finally, we constructed a coarse-grained theory, amenable to analysis, for centrosome motion that directly relates the biophysical properties of MT nucleation, growth, and interaction with cortical force generators, to the cell biological behavior of centrosomes. This theory demonstrates that cortical pulling alone is sufficient to explain centrosome behaviors. Taken together, our results argue that cortical pulling forces provide a unifying explanation of the diverse centrosome motions - directed, stable positioning, and oscillations – found in *C. elegans* embryos. Given the ubiquity of these centrosome behaviors across cell biology, and the proposed role of cortical pulling forces in many different contexts, the investigative framework presented here should be widely applicable.

## RESULTS

### Pulling forces drive spindle oscillations

We began by studying the transverse oscillations of the first mitotic spindle in late metaphase to anaphase in *C. elegans* embryos (Figure 1A). We used spinning disk confocal microscopy to image GFP::β-tubulin and mCherry::γ-tubulin, and tracked the motion of the spindle poles using automated image analysis (see STAR Methods). As others have noted, the anterior pole (Figure 1A,1B orange) and the posterior pole (Figure 1A, 1B blue) oscillate out of phase with each other (Movie S1), with the posterior pole displaying larger amplitudes, and more robust oscillations (Pecreaux et al., 2006, Grill et al., 2005). As described above, the velocity of a centrosome is proportional to the force acting on it. Thus the oscillatory motion of the centrosomes must be driven by oscillations in the magnitudes and directions of the forces exerted on the centrosomes: when a centrosome reverses its motion at the peak of an oscillation, with zero velocity (Figure 1B, insert, T_p_), there is zero net force acting on it; when the centrosome is at the midpoint of an oscillation, moving with maximum speed (Figure 1B, insert, T_m_), then the forces acting on the centrosome are maximal and in the direction of motion.

**Figure 1.**
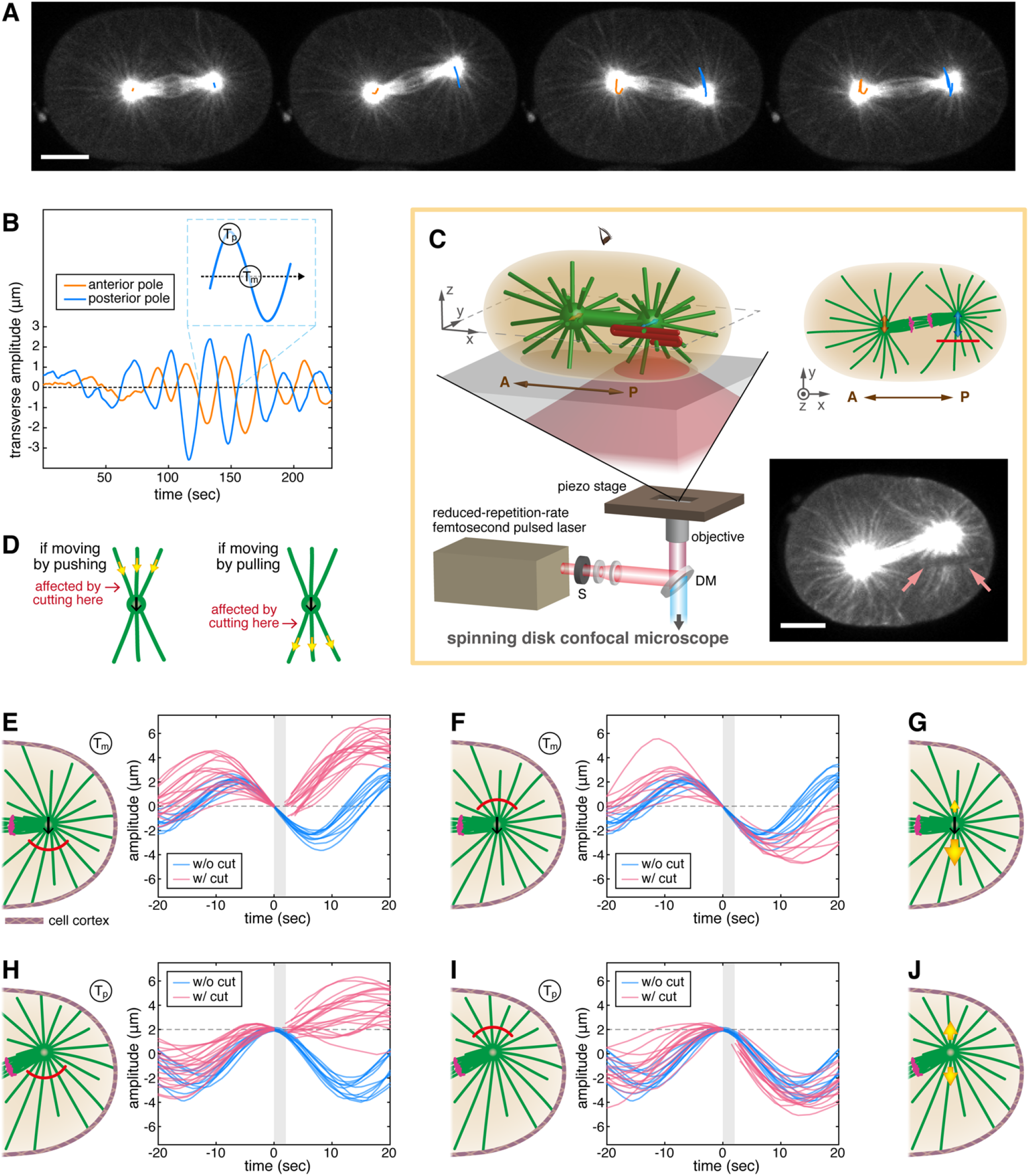
Laser Ablation during Spindle Oscillations in the first mitotic division of *C. elegans*. (A and B): Spindle oscillations in a one-cell *C. elegans* embryo labeled with GFP::β-tubulin. (A) One oscillation cycle and the corresponding trajectories of the anterior (orange) and posterior (blue) centrosomes (scale bar: 10 μm). (B) The transverse amplitudes of the centrosomes and the definition of times T_p_ and T_m_. (C) (left) Femtosecond Stereotactic Laser Ablation (FESLA): the focus of a Ti:sapphire femtosecond pulsed laser with reduced repetition rate (16~80-KHz) is scanned over the sample in a complex 3D pattern (red line) for ablation using a 3-axis piezo stage (microtubules(MTs) in green; S: shutter; DM: dichroic mirror). (right, upper) A 2D view of the x-y imaging plane (magenta, chromosomes; orange arrow, motion of anterior centrosome; blue arrow, motion of posterior centrosome; brown arrow, the anterior(A)-to-posterior(P) axis). (right, lower) Image of a sample after ablation, arrows indicate the cut region (scale bar: 10 μm). (D) Illustration of how cutting at different locations can distinguish between pushing and pulling scenarios. (E-J) Arc cuts ablating astral MTs on either transverse side of posterior centrosomes at two different time points in the oscillation cycle. At time T_m_ (E-G), the arc cuts performed in front of centrosomes (E) and at the rear of centrosomes (F), give the proposed net forces (yellow arrows) (G). At time T_p_ (H-J), the arc cuts carried out below centrosomes (H) and above centrosomes (I), give the proposed net forces (J).

To investigate how astral MTs contribute to these motions, we utilized a novel femtosecond stereotactic laser ablation (FESLA) system capable of cutting 3D patterns with highly controlled timing and location (Figure 1C, see STAR Methods) (Farhadifar et al., 2020, Yu et al., 2019). FESLA utilizes a reduced-repetition-rate ultrafast femtosecond laser to produce highly localized cuts with submicron precision (Chung et al., 2006, Chung and Mazur, 2009, Gabel et al., 2008, Gabel et al., 2007, Vogel et al., 2005). We sought to determine the relative contribution of pushing and pulling forces to spindle oscillations by selectively cutting different populations of astral MTs: if a centrosome is being pushed from the rear, then cutting the astral MTs behind the centrosome will cause it to stop (Figure 1D, left), while if a centrosome is being pulled forward, then cutting astral MTs in the front will halt its motion (Figure 1D, right). We first used this approach to explore the forces acting on centrosomes when they are at the midpoint of their oscillation, moving with maximum speed (Figure 1B, insert, T_m_). When astral MTs in front of these centrosomes were cut, the centrosomes immediately stopped moving along their original course and then rapidly proceeded in the opposite direction (Figure 1E, Movie S2, and Figure S2D left; *v_y_* is negative at T_m_, and *Δv_y_* = *v_y(after)_* - *v_y(before)_*; *Δv_y_* = 0.05 ± 0.05 μm/sec in 11 uncut embryos; while *Δv_y_* = 0.86 ± 0.06 μm/sec in 18 arc-cut embryos; data are shown as mean ± SEM, and *p*=5.0×10^−10^ calculated by two-tailed Student’s *t*-test). As cutting astral MTs in front of these centrosomes stopped their motion, this result strongly argues that this forward movement was primarily driven by net pulling from these astral MTs. Since the centrosomes subsequently moved in the opposite direction, it argues that pulling forces are also exerted by the astral MTs at the rear of the centrosome. Cutting astral MTs at the rear of the centrosome during this same point in the oscillations only marginally impacted their velocity (Figure 1F, Movie S2, and Figure S2D left; *Δv_y_* = 0.05 ± 0.05 μm/sec in 11 uncut embryos; and *Δv_y_* = −0.06 ± 0.04 μm/sec in 11 arc-cut embryos; *p*=0.087), arguing that the downward pulling forces greatly dominate over upward pulling forces. Taken together, these results argue that, at the oscillation midpoints, astral MTs exert net pulling forces on centrosomes, with larger pulling force from the front and smaller pulling force from the rear (Figure 1G).

We next investigated the forces acting on centrosomes at the peak of the oscillation, when their velocity is zero (Figure 1B, insert, T_p_). Cutting the astral MTs below the centrosome, that faced the distant cortex, inhibited the downward motion that occurs subsequent to this time in control embryos (Figure 1H, Movie S2, and Figure S2D right; *Δv_y_* = −0.55 ± 0.04 μm/sec in 11 uncut embryos; while *Δv_y_* = 0.18 ± 0.05 μm/sec in 21 arc-cut embryos; *p*=5.8×10^−10^). After cutting, many of the centrosomes moved in the opposite direction: upward towards the near cortex. As cutting astral MTs below these centrosomes prevented their downward motion, this result strongly argues that the downward movement in control embryos is primarily driven by pulling from these astral MTs. Since the centrosomes subsequently move in the opposite direction, it suggests that pulling forces are also exerted by the astral MTs above the centrosome. Cutting the astral MTs above the centrosome during this same point in the oscillation did not significantly impact the centrosomes subsequent motion (Figure 1I, Movie S2, and Figure S2D right; *Δv_y_* = −0.55 ± 0.04 μm/sec in 11 uncut embryos; and *Δv_y_* = −0.61 ± 0.04 μm/sec in 16 arc-cut embryos; *p*=0.28). Taken together, these results argue that, at the peaks of the oscillations, astral MTs exert net pulling forces on centrosomes. Since the centrosome has zero velocity at this point, the pulling forces from above and below the centrosome must be equal and opposite (Figure 1J). Thus, astral MTs on both transverse sides of the centrosome exert pulling forces throughout the oscillations, with a magnitude and net effect that differs at different points in the oscillation cycle.

We next investigated the contribution of additional populations of astral MTs to spindle oscillations. We performed cuts in the shape of an open cylinder around the posterior centrosome with the cylinder axis aligned with the transverse (y-)axis (Figure 2A), thereby severing MTs perpendicular to the direction of the oscillation (Figure 2B and 2C, and Movie S3). Centrosomes continued to oscillate after these cylindrical cuts (Figure 2D), with an approximately twofold increase in amplitude immediately after the cut (Figure 2E, and Figure S2G and S2H, *A_after_ /A_before_* = 1.6 ± 0.1 in 22 uncut cycles; and *A_after_ /A_before_* = 3.5 ± 0.4 in 15 embryos with open cylinder cuts; *p*=1.8×10^−5^). Since the centrosomes still reverse direction after the cylindrical cut, this result demonstrates that the bending of the MTs perpendicular to the oscillation axis are not required for the restoring mechanism in oscillations (Kozlowski et al., 2007). We next performed a double-(y-z-)plane cut (Figure S2E) ablating MTs in two planes orthogonal to the spindle axis, while leaving the perpendicular MTs of the remaining two planes intact. We found that the more perpendicular MTs that were ablated, the more the oscillation amplitude increased (Figure S2H). This is consistent with the perpendicular MTs being subject to pulling forces, and hence constraining centrosome motion.

**Figure 2.**
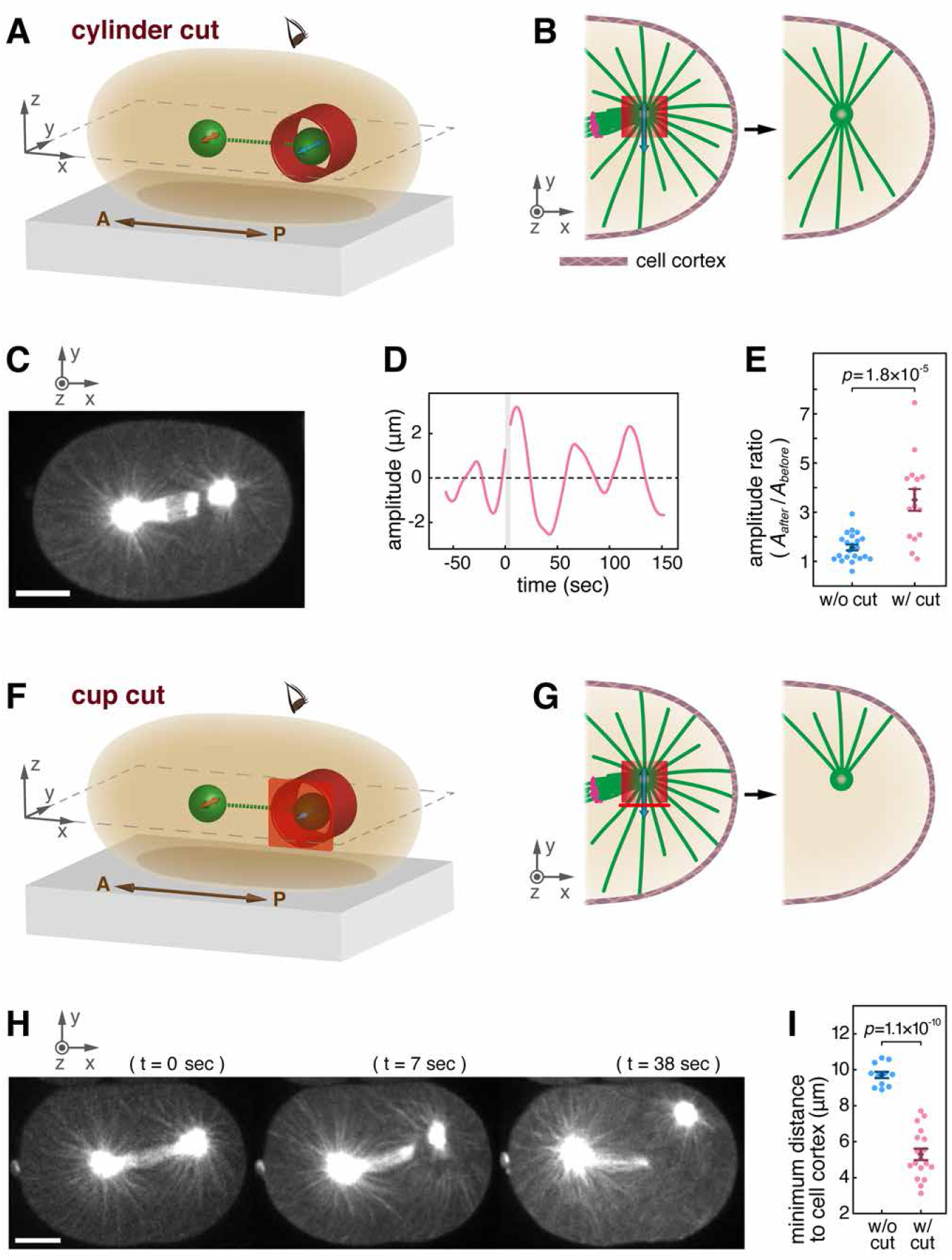
Cylinder Cuts and Cup Cuts Around the Posterior Centrosomes during Spindle Oscillations. Spindle poles are illustrated by green balls connected with green dashed lines (A: anterior, P: posterior) in 3D schematics (A and F). The posterior portions of the x-y mid-planes (view from the top) are shown in the corresponding 2D schematics (B and G), with spindles and astral MTs in green, and chromosomes in magenta. Ablation geometrics are portrayed in red in all schematics. (A-E) Open cylinder cuts: A 3D schematic (A) of the cut aligned along the transverse oscillation (y-)axis, and the 2D schematic (B) showing the ablation of the MTs perpendicular to the oscillation, leaving the transverse astral MTs intact. (C) An image taken directly after a cylinder cut (scale bar: 10 μm). (D) An example of oscillation amplitude before and after the cut, with the time of ablation marked by the light gray vertical strip. (E) The amplitude of oscillations increases after cylinder cuts. (F-I) Cup cuts: A 3D schematic (F) of the cut whose opening facing in the transverse direction, and the 2D schematic (G) showing the ablation of all astral MTs except those extending out from the opening of the cup. (H) Images before and after a cup cut (scale bar: 10 μm). (I) The comparison of the minimum distances, from approaching posterior centrosomes to cell cortices, between uncut and cup-cut embryos (see Figure S2J for distance measurement). The *p*-values are calculated using two-tailed Student’s *t*-test. (Error bars: SEM)

We next studied the role of MT pushing. It is possible that substantial pulling and pushing forces can simultaneously be exerted by different astral MTs located on the same side of the centrosome. To test this, we performed cup-shape cuts to temporarily eliminate all MTs around centrosomes except for those extending to one transverse side (Figure 2F and 2G). We observed that the centrosome rapidly moved very close to the cortex (Figure 2H and Movie S3) after a cup cut, approaching a minimum distance of 5.3 ± 0.3 μm (n=18) from the cortex, compared to a distance of 9.7 ± 0.2 μm (n=11) in uncut embryos (Figure 2I; *p*=1.1×10^−10^). Thus, pulling forces dominate over pushing forces from astral MTs on the same side of a centrosome. Furthermore, the observation that centrosomes approach very close to the cortices after cup cuts argues that pushing forces do not significantly contribute to the restoring force during normal oscillations.

### During transverse oscillations, pulling forces result from force generators on the cell cortex

We next investigated if pulling forces that act on centrosomes during the late metaphase to anaphase transverse oscillations result from force generators in the cytoplasm or on the cell cortex (Figure 3A). For force generators in the cytoplasm, such as from dynein transporting organelles along MTs (Kimura and Onami, 2005, Kimura and Kimura, 2011, Shinar et al., 2011, Xie and Minc, 2020), the pulling forces on astral MTs are balanced by equal and opposite forces on the cytoplasm. Due to the viscous nature of the cytoplasm (Daniels et al., 2006, Wirtz, 2009), these forces acting on the cytoplasm will generate flows, which will tend to be in the direction opposite MT motion (Figure 3A, top). For force generators anchored to the cell cortex, pulling forces on astral MTs are balanced by equal and opposite forces on the cell cortex (Figure 3A, bottom). In this case, as an MT moves it will drag fluid (i.e. the cytoplasm) along with it, so that cytoplasmic flows and MT motions will tend to be in the same direction. This scenario holds irrespective of whether the cortically anchored pulling forces are generated by dynein, MT depolymerization, or some other mechanism (Cowan and Hyman, 2004, Kotak et al., 2012, Nguyen-Ngoc et al., 2007, Gusnowski and Srayko, 2011, Laan et al., 2012). Since cytoplasmically based pulling forces tend to produce flows in the direction opposite to MT motions, while cortically based pulling forces tend to produce flows in the same direction as MT motions, measuring fluid flow provides a means to distinguish these two possibilities. However, the detailed pattern of flows in an embryo can be difficult to predict because of the long-range nature of hydrodynamic interactions due to flow incompressiblity, and the effects of the cell boundary. We thus turned to large scale computer simulations to study the pattern of flows for cytoplasmically and cortically based pulling forces (Nazockdast et al., 2017a, Nazockdast et al., 2017c) (see STAR Methods).

**Figure 3.**
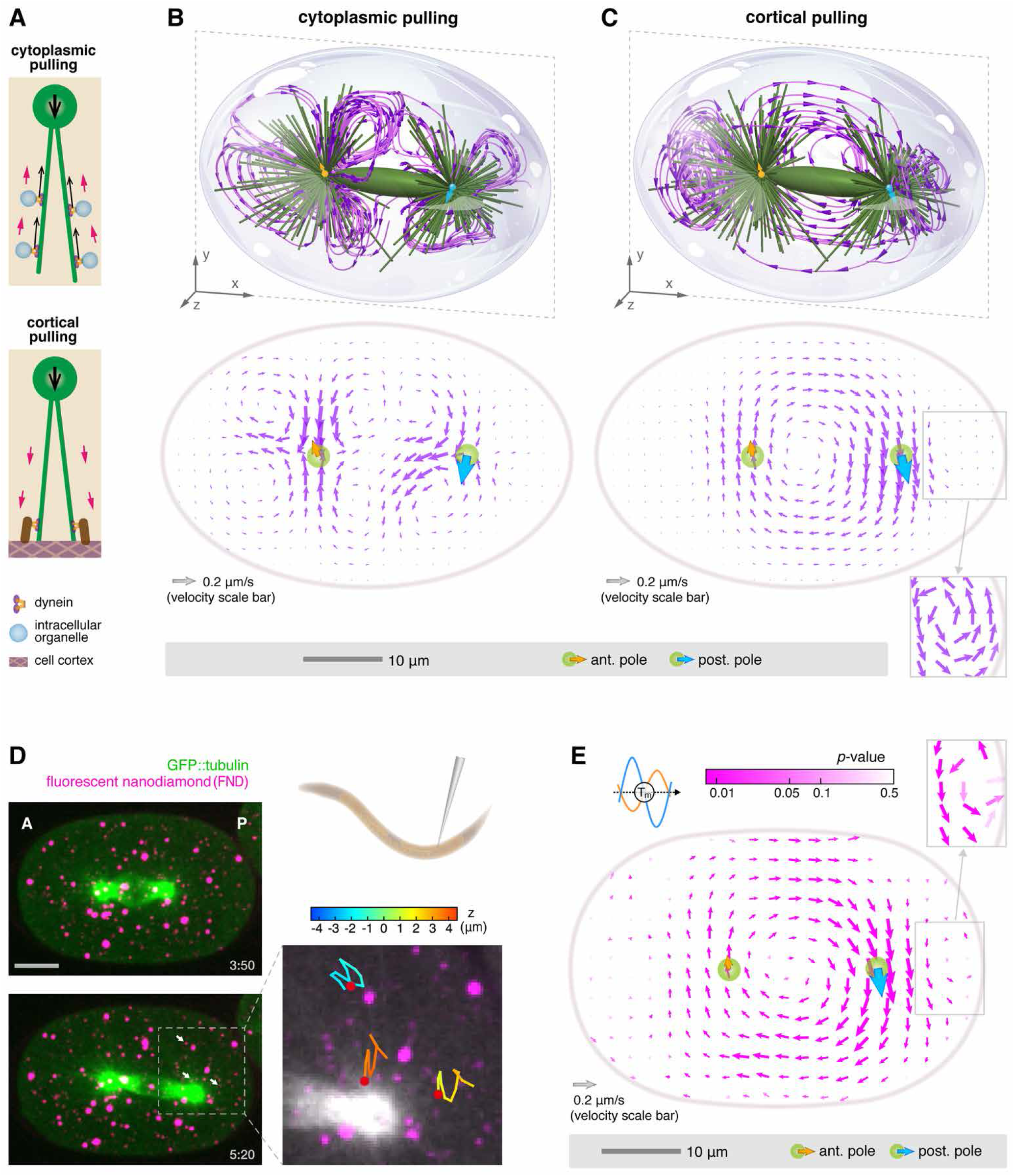
Fluid Flows during Spindle Oscillations. (A) Schematic of fluid flow directions near a centrosome (green circle) and astral MTs (green lines). (top) In a cytoplasmic pulling model, the direction of fluid flow (magenta arrows) is opposite that of centrosome motion (bold black arrow). (bottom) In a cortical pulling model, fluid flow and centrosome motion are in the same direction. (B and C) Top panels: 3D computational fluid dynamics simulations near the midpoint of oscillations, time T_m_, with cytoplasmic pulling (B) or cortical pulling (C). Bottom panels: The averaged 2D-projection fluid vector fields (in the x-y plane) from the above simulations. (D) (left) Two time frames (Δt = 90 sec), showing maximal intensity z-projection of microinjected (upper right) fluorescent nanodiamonds (FNDs). (lower right) Illustrative 50-sec 3D trajectories from 3 FNDs (identified by the white arrows, lower left) (Scale bar: 10 μm; A: anterior, P: posterior) (E) Experimentally measured fluid flow vector field near the oscillation midpoint, time T_m_, obtained by tracking FNDs from 22 embryos and averaging their projected x-y velocities. The length of the arrows is proportional to the flow velocity. The intensity of the arrows indicates the statistical significance of the flow speed (*p*-value colormap) (see STAR Methods for details). Arrows on centrosomes indicate mean measured centrosome velocities. The zoom-in backflows in (C) and (E) are displayed with fixed-length vectors.

We first simulated the effect of cytoplasmic forces generated by dynein transporting cargo along MTs at time T_m_ (Figure 1B, insert). These simulations show that a complex, 3D flow results, consisting of multiple vortices (Figure 3B, top). To help visualize this flow, we calculated the 2D projection of this flow in the plane of spindle motion (Figure 3B, bottom). This flow pattern occurs because cytoplasmic pulling forces tend to create minus-end directed flows along MTs, but the incompressibility of the fluid prevents these flows from being uniformly directed inward toward centrosomes. A very different pattern of flow is produced by simulating cortical pulling forces, as seen in both 3D (Figure 3C, top) or in the 2D projection in the plane of spindle motion (Figure 3C, bottom). In this case, the downward motion of the posterior centrosome leads to an overall rotational flow, with flow speeds of the same scale as the spindle poles’ speeds. A much smaller, counter-rotating vortex is also produced between the posterior centrosome and the cortex (Figure 3C, bottom insert).

With these predictions in hand, we next sought to experimentally measure cytoplasmic fluid flows in *C. elegans* embryos during spindle oscillations. To this end, we microinjected passivated (Daniels et al., 2006, Valentine et al., 2004) fluorescent nanodiamonds (FNDs) (Mochalin et al., 2012, Fu et al., 2007, Mohan et al., 2010, Vaijayanthimala et al., 2012, Chang et al., 2008, Su et al., 2017) into *C. elegans* syncytial gonads, which subsequently became incorporated into embryos (Figure 3D, see STAR Methods). We used 3D time-lapse spinning disk confocal microscopy to image the FNDs and the spindle, which were visualized by GFP::β-tubulin (Figure 3D and Movie S4). We tracked the FNDs in 3D using automated image analysis to find their trajectories over time (Figure 3D right). We averaged data from multiple embryos (n=22) and multiple oscillations to better reveal the underlying fluid flow (see Figure S3A and S3B and STAR Methods). Figure 3E shows the resulting measured fluid flow throughout the embryo when the centrosomes are at the midpoint of their oscillations and moving with maximum speed (Figure 1B, insert, T_m_). The measured fluid flow is remarkably similar to that calculated from the cortical pulling model (compare Figure 3C and Figure 3E), displaying both the characteristic circular motions around the two spindle poles, as well as a subtler counter-rotating backflow between the posterior centrosome and the cortex (see both inserts in Figure 3C and Figure 3E). Also like the simulations for cortical pulling, the measured fluid speeds are similar in magnitude to the measured pole speeds. If both cytoplasmic and cortical pulling forces were present, then the resulting flows, and centrosome velocities, would be a linear combination of those shown in Figures 3B and 3C (see STAR Methods). Since, within experimental error, the measured flow agrees with the pattern observed in simulations of cortical pulling, with no sign of the complex flows predicted from cytoplasmic pulling, we estimate that the contributions to the centrosomes velocities from cortical pulling forces are many-fold greater than those from cytoplasmic pulling forces. In sum, these results strongly argue that the pulling forces acting on centrosomes during anaphase spindle oscillations are predominately cortically based.

### In prometaphase and metaphase, cortical pulling forces stably position the spindle

During the first mitotic division of *C. elegans*, the spindle forms near the cell center and is aligned along the cell’s long axis. The spindle remains centered in prometaphase and metaphase, and slowly elongates before entering anaphase. We next investigated the nature of the forces acting on the spindle at these times.

We first used FESLA to test if astral MTs exert pushing or pulling forces on the spindle when it is stably centered. We separately probed astral MTs transverse and longitudinal to the spindle axis. Ablating 9~11-μm(x) by 6-μm(z) planes 2.5~5 μm below the posterior centrosomes caused the centrosomes to immediately displace upward, away from the cut astral MTs, arguing that these MTs exerted net pulling forces on the centrosomes (Figure 4A and Movie S5). Similarly, ablating 10.5~11.5-μm(y) by 6-μm(z) planes located 2.5~5 μm posterior to the posterior centrosome caused the spindles to immediately displace away from the cut astral MTs, arguing that these MTs also exert net pulling forces on the spindles (Figure 4B and Movie S5). After both sets of cuts, the astral MTs recovered in approximately 30 seconds and the centrosomes returned to their original positions. These results imply that the position of the spindle is actively maintained by the balance of astral MTs exerting net pulling forces from different directions.

**Figure 4.**
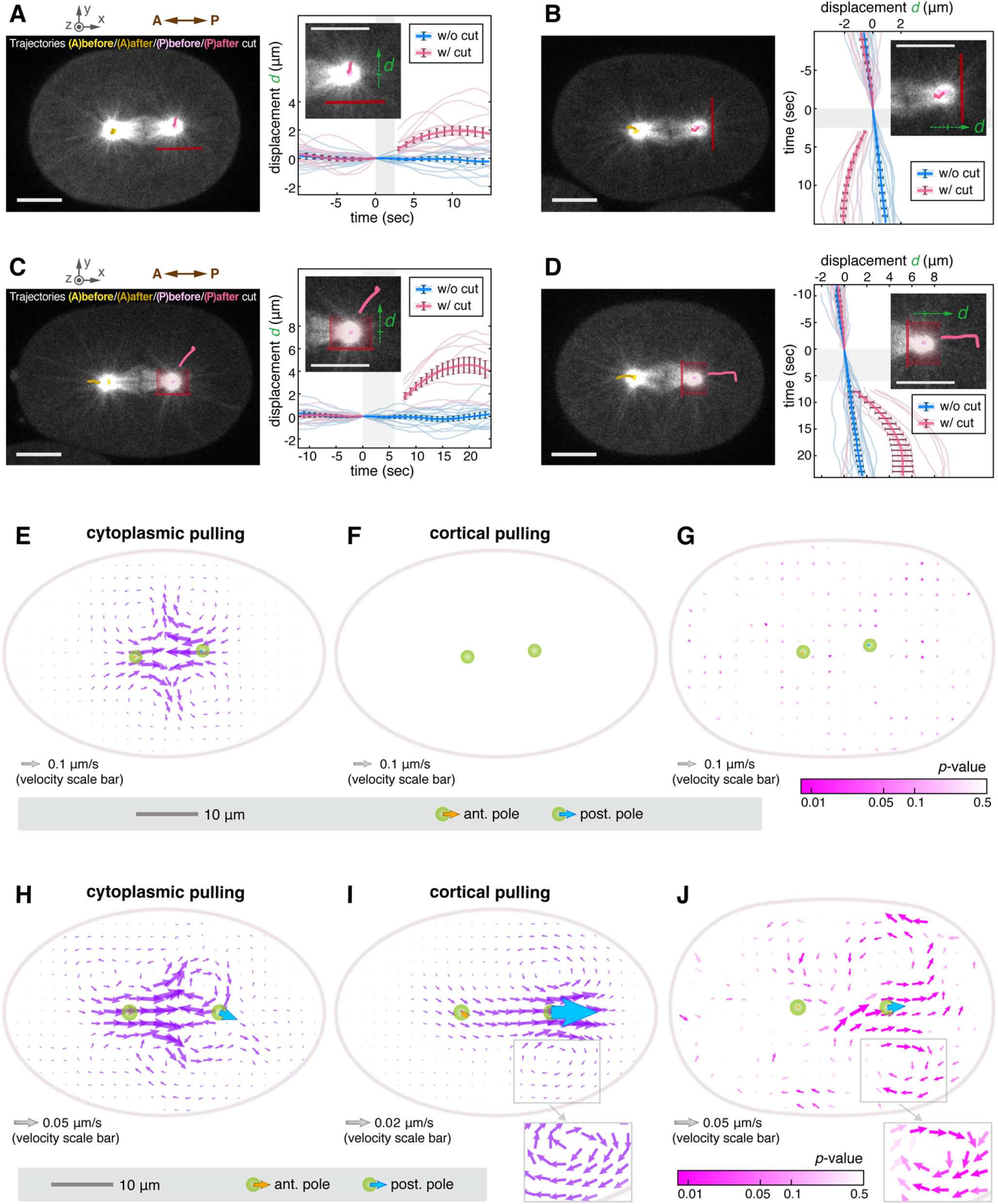
Laser Ablation and Fluid Flows in Prometaphase and Metaphase. (A-D) Laser ablation performed in metaphase (cutting patterns in red). The centrosome trajectories before and after cutting are plotted on the images with the designated colors. Right panels illustrate the definition of displacement *d* for the corresponding cuts in their inserted images, and show the centrosome displacement data for uncut and cut embryos with the mean value curves and SEM error bars overlaying the semi-transparent raw data curves. (Scale bar: 10 μm; A: anterior, P: posterior) (A) The (x-z-)plane cuts on the transverse sides of the posterior centrosomes. (B) The (y-z-)plane cuts located posteriorly to the posterior centrosomes. (C and D) The cup cuts surrounding the posterior centrosomes with the opening mouths facing the transverse sides (C) or the posterior ends (D). (E and F) The simulated fluid flows in prometaphase under (E) cytoplasmic pulling or (F) cortical pulling models. (G) The flow vector field of averaged experimental results from tracking FNDs in 9 embryos in prometaphase. (H and I) The simulated fluid flows during metaphase spindle elongation under (H) cytoplasmic pulling or (I) cortical pulling models. (J) The experimental flow vector field derived from averaging the movements of tracked FNDs in 14 embryos during metaphase spindle elongation. The zoom-in highlighted backflows in (I) and (J) are exhibited with fixed-length vectors. The statistical significance of the velocity vectors is indicated by their color intensity (*p*-value colormap) in (G) and (J).

To determine if the net pulling force results from a combination of both pulling and pushing forces from different astral MTs on the same side of a centrosome, we used cup cuts as described above. We performed cup cuts, leaving only a cone of astral MTs associated with the centrosome, emanating in either the transverse (Figure 4C and Movie S5) or longitudinal (Figure 4D and Movie S5) directions. In both cases, centrosomes rapidly moved in the direction of the remaining astral MTs, displaced farther than in response to plane cuts, and only slowed down once astral MTs in the ablated regions began to recover (Movie S5). Thus, these experiments provide no sign of MT pushing forces, even when the centrosomes are displaced by ~5 μm, suggesting a minimal contribution of pushing to the forces that position the spindle near the cell center.

The pulling forces that stably center the spindle in prometaphase and metaphase could originate either in the cytoplasm or at the cortex. We again used a combination of large-scale fluid dynamics simulations and measurements of fluid flow to distinguish these possibilities. Simulations of cytoplasmic pulling display extensive flows, organized into vortices (Figure 4E). Such large flows are necessarily present in a cytoplasmic pulling model, even when the spindle is stationary, as they are ultimately responsible for the forces that maintain the position of the spindle in this model. In contrast, flows only result from the motion of the spindle in a cortical pulling model, so are absent when the spindle is stationary (Figure 4F). To experimentally measure fluid flow, we tracked FNDs in 9 embryos in prometaphase, during which the spindle displayed no appreciable motion, and averaged the results together. No coherent fluid motion was present (Figure 4G), consistent with predictions from the cortical pulling model and inconsistent with cytoplasmic pulling. We next investigated fluid flow during the slow elongation of the spindle in metaphase, before the onset of (late metaphase to anaphase) spindle oscillations, when the posterior centrosome moves and the anterior centrosome is mostly stationary. Simulations of the cytoplasmic pulling model again produce complex flows (Figure 4H), including two counter-rotating vortices in the posterior half, with fluid flow locally moving in the direction opposite to the posterior centrosome. Simulations of the cortical pulling model also produce vortices in the posterior half of the cell, but in this case the fluid flow locally moves in the same direction as the posterior centrosome (Figure 4I and the insert). Averaging together the trajectories of FNDs from 14 embryos with their spindles elongating in metaphase resulted in fluid flow with a pattern and amplitude quite similar to those predicted by cortical pulling, and inconsistent with the cytoplasmic pulling model (Figure 4J and the insert). As described above, if cytoplasmic and cortical pulling forces were simultaneously acting, then the resulting flows, and centrosome velocities, would be a linear combination of the predicted flows of those forces acting independently of each other (see STAR Methods). Since, within experimental error, the measured flow agrees with the pattern observed in simulations of cortical pulling, we estimate that the contributions to velocities from cortical pulling forces are many-fold greater than those from cytoplasmic pulling forces.

Taken together, through FESLA and by comparing fluid flow measurements with large-scale fluid dynamics simulations, we demonstrated that spindle positioning in prometaphase and metaphase is primarily driven by cortical pulling forces, rather than from MT pushing or cytoplasmic pulling.

### Cortical pulling forces drive pronuclear migration/centration and rotation

At an even earlier stage, the female and male pronuclei meet near the posterior cortex, and the pronuclear complex (PNC) migrates toward the cell center. During the centration process, the PNC rotates to align with the long axis of the cell (Coffman et al., 2016). We next investigated the nature of the forces acting on the PNC at these times.

We first used FESLA to test if astral MTs exert pushing or pulling forces on the PNC during centration and rotation. We separately probed astral MTs associated with the leading or trailing centrosomes by ablating 11~15-μm(x-y) by 6-μm(z) planes 2~4 μm microns away from the centrosome (Figure 5A and Movie S6). In both cases, the centrosomes rapidly moved away from the cut astral MTs (Figure S4F and S4G), which is clearly seen by realigning the centrosomes relative to the cut (Figure 5B). These results imply that astral MTs primarily exert pulling forces on centrosomes during pronuclear centration.

**Figure 5.**
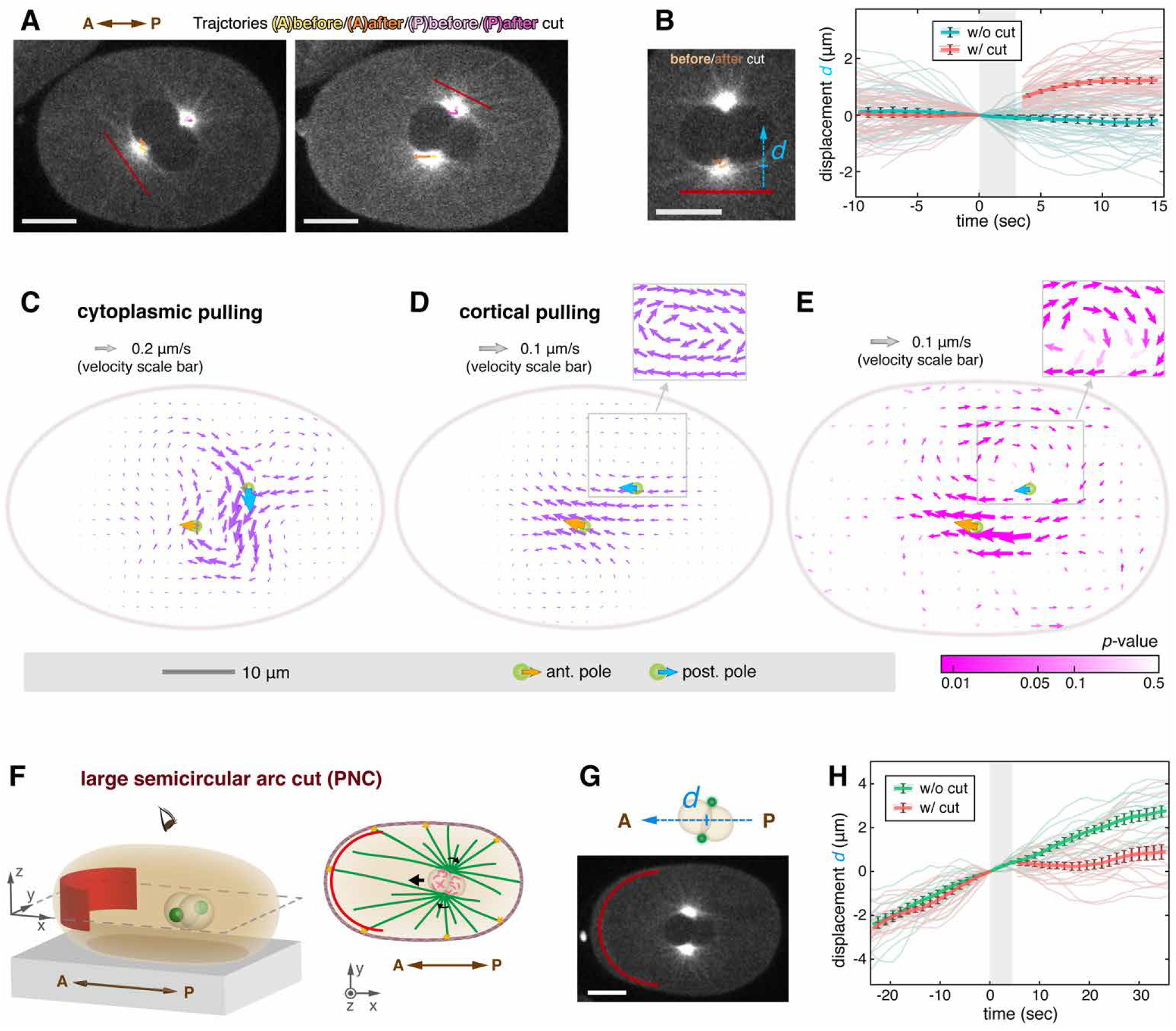
Laser Ablation and Fluid Flows during Migration of the Pronuclear Complex (PNC) (A and B) The plane cuts (ablation drawn in red) during PNC migration. (A) Images indicating the locations of plane cuts (left: cutting near the anterior or leading centrosome; right: cutting near the posterior or trailing one). On both images, the centrosome trajectories within the windows of 10 sec before and 15 sec after cutting are plotted. (A: anterior or leading, P: posterior or trailing) (B) The definition of centrosome displacement *d*, the relative orthogonal (away from the cut) displacement from the locus right before cutting, is illustrated on the left. The right panel presents the uncut and cut displacement data sets. (C and D) The simulated fluid flows under (C) cytoplasmic pulling or (D) cortical pulling models. (E) The experimental flow vector field obtained from averaging 8 embryos with tracked FNDs. The statistical significance of the velocity vectors is indicated by their color (p-value colormap). (F-H) Large semicircular arc cuts close to the anterior cortices during PNC migration. (F) The 3D schematic (left) and the 2D schematic (right) of the arc cut (drawn in red). The black arrows labeled on the 2D schematic indicates PNC translation and rotation. (A: anterior, P: posterior) (G) An image taken right after a semicircular arc cut. To quantify the progression of the PNC migration, the displacement *d* is calculated by projecting the PNC center onto the embryo long axis. (H) The migration displacements of uncut and cut embryos. For each ablated embryo, the zero point (*d* = 0) is defined by the PNC location directly before cutting. All scale bars in images: 10 μm. All displacement data are presented with mean value curves and SEM error bars overlaying the semi-transparent raw data curves. In the zoom-in inserts of (D) and (E), the highlighted circular flow patterns are displayed with fixed-length vectors.

We next studied fluid flow to investigate if the pulling forces on the pronuclei originate in the cytoplasm or at the cortex. Simulations of cytoplasmic pulling during rotation and centration produce a complex pattern of flows (Figure 5C). In contrast, flows only result from, and thus locally follow, the motion of the PNC in a cortical pulling model. Consistent with this expectation, simulations of the cortical pulling model produce flows which rotate at the speed of the centrosomes, along with backflows due to the cell boundary and incompressibility of the fluid (Figure 5D and the insert). We measured fluid flows in 8 embryos by tracking FNDs during similar stages of pronuclear migration and rotation and averaging the results together (see STAR Methods). The experimentally determined fluid flow has a similar pattern (Figure 5E and the insert) and amplitude to those predicted by cortical pulling, and is inconsistent with the cytoplasmic pulling model.

Fluid flow measurements support cortical pulling and argue against cytoplasmic pulling, but MTs connecting the PNC to the anterior cortex cannot be clearly visualized (Kimura and Kimura, 2011), and such MTs are necessary to support cortical pulling during pronuclear centration. We thus sought an additional test of the importance of cortical pulling. We performed a large semicircular arc cut in front of the migrating PNC, close to the anterior cortex, such that only MTs a few microns away from the cortex would be ablated (Figure 5F and 5G). The PNC immediately ceased advancing after the cut was performed (Figure 5H and Movie S6), arguing that centration requires astral MTs to contact the anterior cortex. This result is not easily explained by a cytoplasmic pulling model but is expected in a cortical pulling model. In sum, through FESLA and fluid flow analysis, we demonstrated that PNC migration is primarily driven by cortical pulling forces, rather than from MT pushing or cytoplasmic pulling.

### Cortical pulling forces are sufficient to account for centrosome oscillations, stable positioning, and centration

Our above work shows that pulling forces always locally dominate (from FESLA) and these pulling forces result from cortical force generators (by comparing fluid flow measurements with large-scale fluid dynamics simulations) at all times in one-cell *C. elegans* embryos: during anaphase spindle oscillations, during metaphase/prometaphase stable positioning, and during PNC centration. We next investigated if cortical pulling forces alone are sufficient to account for those diverse centrosome motions. We developed a coarse-grained model derived from the biophysics of MT nucleation, growth, and interaction with pulling cortical force generators (see STAR Methods). In this model (Figure 6A), a centrosome nucleates MTs at a rate *γ*, which grow from their plus-end with velocity *V_g_*, and undergo catastrophe (i.e. switch from growing to shrinking) with rate *λ*. MTs that hit an unoccupied force generator bind to it and experience a pulling force *f*_0_, which is transmitted to the centrosome. The force generators are stoichiometric: each force generator can bind to at most one MT at a time (Farhadifar et al., 2020). Bound MTs detach from force generators at a rate *κ*, whereupon they undergo catastrophe. MTs that hit the cell cortex without binding to a force generator undergo immediate catastrophe. We consider a single centrosome moving along the *Y* axis in a spherical cell of radius *W*, uniformly covered with *M* cortical force generators (Figure 6B). The net pulling force on the centrosome is the sum of pulling forces on all its MTs. This net pulling force is balanced by a drag force proportional to the centrosome’s velocity (see STAR Methods).

**Figure 6.**
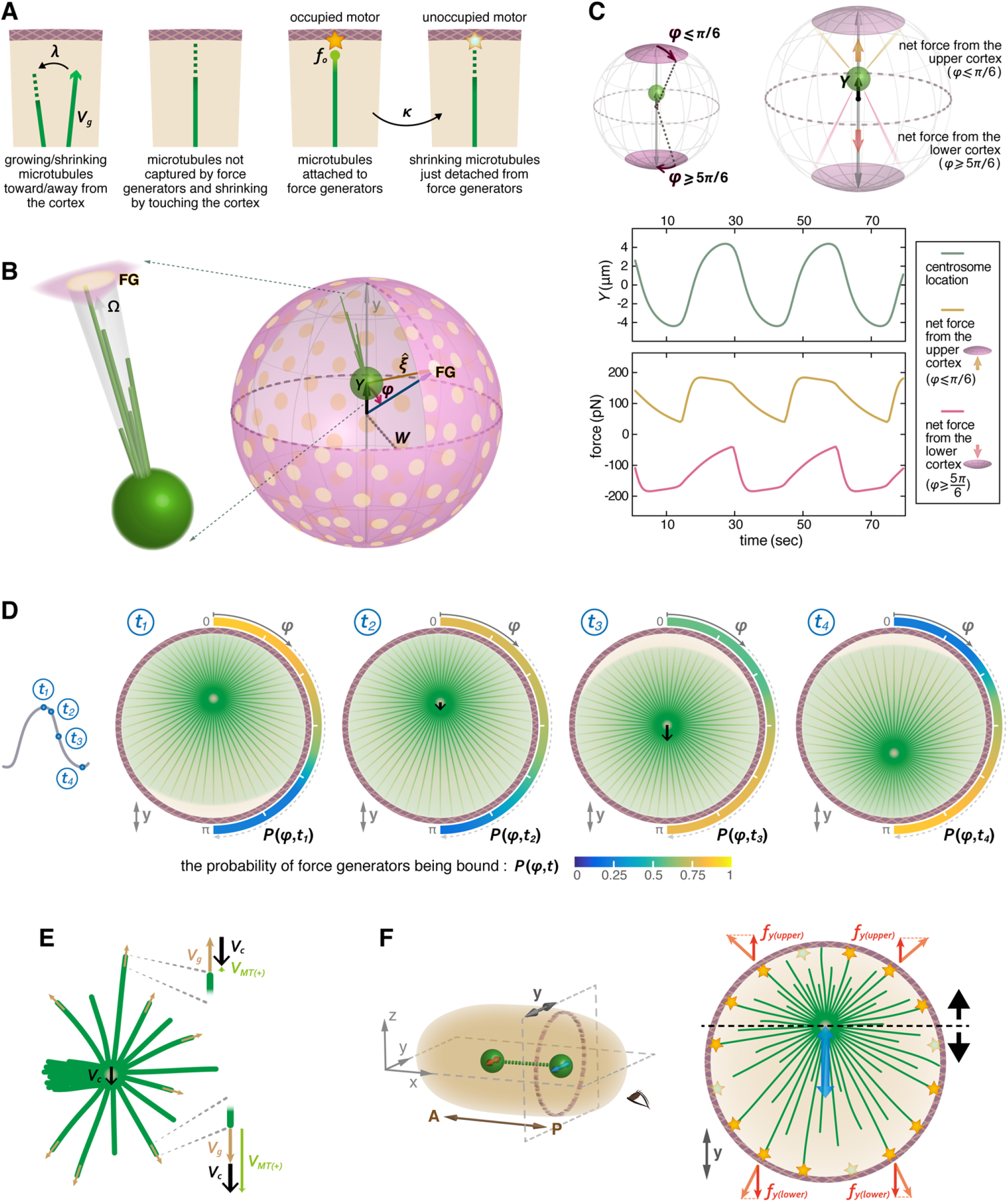
Coarse-grained Model of Cortical Pulling. (A) Key processes in the coarse-grained model. From left to right: MTs grow from their plusends with velocity *V_g_*, and undergo catastrophe (i.e. switch from growing to shrinking) with rate *λ*. MTs that hit the cell cortex without binding to a force generator undergo immediate catastrophe. MTs that hit an unoccupied force generator bind to it and experience a pulling force *f*_0_. Bound MTs detach from force generators at a rate *κ*, whereupon they undergo catastrophe. (B) The geometry of the coarse-grained model. The cell is treated as a sphere of radius *W* decorated with force generators (FG, yellow discs). The centrosome (green sphere) is located at a distance *Y* (along the y-axis) from the center of the cell. 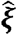 is a unit vector extending from the centrosome to an FG. Ω is the impingement rate of MTs onto a FG. (C) Upper schematic defining the polar caps over which polar pulling forces are measured. Plots show simulated results from the coarse-grained model of oscillation amplitude and net cortical forces from the upper and lower cortices (polar caps) over time. (D) From a simulation of oscillations from the coarse-grained model, the probability of cortical force generators being bound: *P*(*φ*, *t*), a function of polar angle *φ* from the y-axis and time, as displayed by the color scale from the bottom color bar. The information is expressed on a crosssection through the sphere containing the y-axis. The black arrows drawn on centrosomes indicate their velocities; the green field shows the centrosomal MT directions, and the front of unattached plus-ends.. (E) The relations between the centrosome moving speed *V_c_*, the MT growth (polymerization) speed *V_g_*, and the speeds of MT plus-ends *V*_*MT*(+)_. (F) A schematic of the geometric effects leading to the restoring mechanism during oscillation. Viewing from the embryo posterior end, the centrosome is shown to oscillate along the y-direction on the y-z plane (right panel). As the centrosome approaches the upper surface, the force from FGs above the centrosome become more oblique, leading to a smaller upwards force projected along the y-axis.

We first tested if this model, which only contains cortical pulling forces, is sufficient to account for spindle oscillations. We numerically simulated the model, using realistic parameters for anaphase, and found that the centrosome oscillates with a similar amplitude and frequency to those seen experimentally (Figure 6C). Therefore, a model that only contains cortical pulling forces (and no MT bending or pushing forces) is sufficient to explain the experimentally observed oscillations.

We next sought to better understand how a model based solely on cortical pulling forces can produce centrosome oscillations. We used a linear stability analysis to analytically calculate that a centrosome at the center can lose stability to an oscillatory state via a Hopf bifurcation if there is more than a critical number of cortical force generators, given by (see STAR Methods):

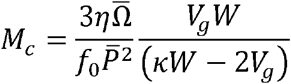

where the steady-state rate of impingement of MTs onto force generators for a stationary centrosome at the cell center before the start of oscillations is 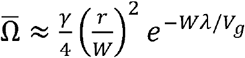, and *r* is the capture radius of a force generator (which includes the distance over which a MT explores the cortex before binding). Here 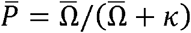 is the steady-state probability of a force generator being bound. In this model, oscillations are only possible if 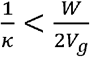, i.e. if the time a MT stays attached to a force generator is sufficiently small compared to the time it takes for a MT to grow across the cell. Thus, the centrosome is driven to oscillate if pulling forces are sufficiently strong and the detachment of MTs from force generators is sufficiently fast.

To gain further insight into the mechanism of oscillations, we returned to numerical simulations and investigated the forces acting on the centrosome. When the centrosome is at the top of the oscillations, the downward pulling force rapidly increases in amplitude, i.e., becomes more negative (Figure 6C). As the centrosome moves toward the lower surface, the downward pulling force remains relatively constant, while the upward pulling force gradually decreases in amplitude. Then, when the centrosome is at the lowest part of the oscillation, the upward pulling force rapidly increases in amplitude and the cycle reverses. The mechanism behind this oscillation in forces is revealed by examining the attachment of MTs to force generators during the oscillations (Figure 6D). When the centrosome is at the top of the oscillations (Figure 6D, *t*_1_), MTs are attached to force generators directly above the centrosome but force generators directly below the centrosome are largely unoccupied. MTs then start to attach to force generators below the centrosome, causing it to start moving downward (Figure 6D, *t*_2_). More MTs continue to attach to the lower side and detach from the upper side (Figure 6D, *t*_3_), and eventually, the process reverses (Figure 6D, *t*_4_).

These calculations and simulations point to an intuitive picture for how oscillations can occur with cortical pulling forces (even when MT pushing or bending forces are absent): during the oscillations, the speed of centrosome motion, ~0.5 μm/sec, is on the order of the MT polymerization velocity, *V_g_* = 0.5 μm/sec. The velocity of a growing MT plus-end is a sum of the MT’s polymerization velocity in the direction of growth and the centrosome velocity (Figure 6E). Thus, the motion of a centrosome away from a force generator reduces the rate of MT impingement on that force generator, while motion towards a force generator increases the rate of impingement (Kozlowski et al., 2007). This results in a decreased probability of attachment, and hence a decreased force, from force generators behind the centrosome, and an increased probability of attachment, and thus an increased force, in front of the centrosome. While this would appear to be a self-reinforcing process, the centrosome speed does eventually reduce. The reduction in speed is caused by a geometric effect: when the centrosome is closer to a surface the force generators pulling it onwards do so from increasingly oblique angles, decreasing their pulling efficacy (Figure 6F). Once the centrosome speed is slowed down, MTs attach to the force generators on the other side of the centrosome, leading to a larger restoring force (because of the geometric effect), and hence oscillations. This phenomenology is robust to details of the system’s geometry: the coarse-grained model also predicts centrosome oscillations can occur between two flat plates (data not shown).

These results demonstrate that theoretically, cortical pulling forces alone are sufficient to produce centrosome oscillations. We next sought to further test this explanation of anaphase spindle oscillations by investigating additional predictions from this model. One key assumption of the model is that MTs that contact the cortex undergo catastrophe and depolymerize. Because of this effect, the simulations predict that the density of (transverse) MTs between the centrosome and the cortex oscillates out of phase with the centrosome’s position relative to the anterior-posterior axis (Figure 7A). We investigated if a similar phenomenon occurs in experiments by measuring the intensity of astral MTs in annular sectors (60°, 3-μm inner radii, 9-μm outer radii) transverse to the spindle oscillations (Figure 7B and Movie S7). The density of transverse MTs oscillates out of phase with centrosome’s position (Figure 7B) in a manner highly reminiscent of the theoretical prediction (Figure 7A). The agreement between the model and experiments for the oscillation in density of astral MTs supports the contention that MT depolymerization is induced upon contact with the cortex (Kozlowski et al., 2007).

**Figure 7.**
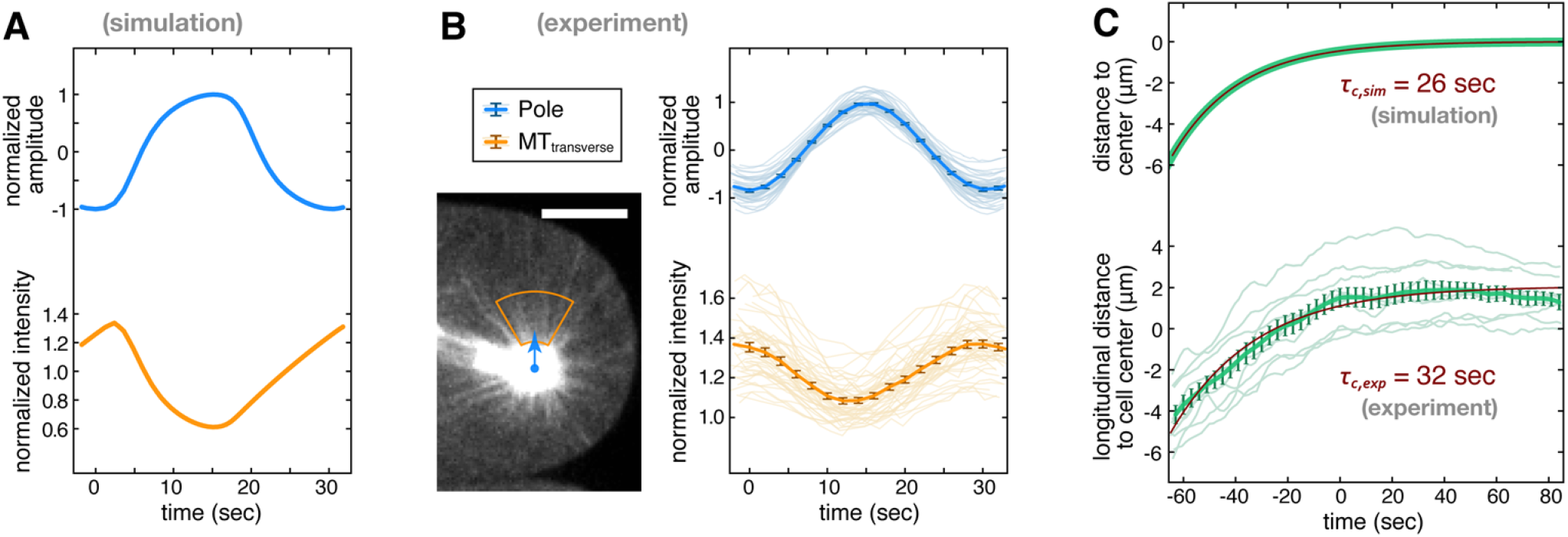
Theory Predictions and Experimental Measurements on Astral MT Density and Pronuclear Centration. (A) Simulation predicts that the density of astral MTs (lower) in an annular sector transverse to the movement of the centrosome oscillates out of phase with the position of the centrosome (upper). (B) Experimental measurement of the intensity of astral MTs in an annular sector transverse to the movement of the centrosome (scale bar: 10 μm). (C) The simulated and experimental results of pronuclear centration and with exponential fits and resulting centering time scales. All experimental data are presented with mean value curves and SEM error bars overlaying the semi-transparent raw data curves.

We next investigated if cortical pulling forces are sufficient to explain the stable positioning of the spindle near the cell center in prometaphase and metaphase. As described above, centrosome oscillations can be accounted for by a model that only contains cortical pulling. If the number of cortical force generators is reduced sufficiently in this model, then the centrosome ceases to oscillate and is stably centered (see STAR Methods). In this model, cortical pulling forces stably position centrosomes because of two effects: 1) the geometric effect described above (Figure 6F), in which the net pulling forces from a surface are reduced as a centrosome approaches it due to the increasingly oblique angles between the axis of displacement and force generators; 2) the stoichiometric interactions between MTs and force generators: i.e. each force generator can bind to at most one MT at a time (Farhadifar et al., 2020). In the absence of stoichiometric interactions, cortical pulling forces are always destabilizing and can never stably center centrosomes (see STAR Methods).

When the centrosome is stably centered, as in prometaphase and metaphase, the resulting spring constant can be analytically calculated as (see STAR Methods):

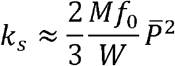

where, as above, *M* is the number of force generators, *f*_0_ is the pulling force exerted by a force generator on a bound MT, 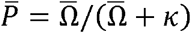 is the steady-state probability of a force generator being bound, and *κ* is the rate at which bound MTs detach from force generators. Using the same parameters to those that reproduced anaphase spindle oscillations in the stochastic simulations, but now only lowering *M*, the number of cortical force generators, this simulation gives a spring constant for spindle centering of 4.8 pN/μm, which is the same order of magnitude as the measurements of Garzon-Coral *et al* (16.4 ± 2.1 pN/μm) (Garzon-Coral et al., 2016). Therefore, a model with only cortical pulling forces is sufficient to explain the stable centering of the spindle in prometaphase and metaphase.

We next investigated if cortical pulling forces are sufficient to explain the PNC centration process. We used the same cortical pulling model which reproduced anaphase oscillations and stable centering in prometaphase and metaphase, and assumed the same number of cortical force generators as in prometaphase and metaphase. The coarse-grained theory can be used to analytically calculate (see STAR Methods) that the PNC should exponentially approach the center with a time scale, *τ_c_*, given by:

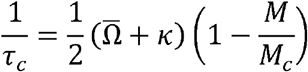

where, as above, 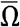 is the steady-state rate that growing MTs impinge upon force generators, *κ* is the rate bound MTs detach from force generators, *M* is the number of motors, and *M_c_* is the critical number of motors that lead to oscillations (as mentioned before, *M* < *M_c_* at these times). Consistent with this prediction, exponential relaxation is observed in both simulations and experiments (Figure 7C), with time-scales of 26 sec and 32 sec (95% confidence interval: 29-36sec) respectively. A second prediction of the coarse-grained theory is that the centering time scale, *τ_c_*, is the same as the time scale, *τ_f_*, of approach to a new equilibrium in response to an applied force. Consistent with this prediction, the observed centering times quoted above are the same as the force response time scale found in simulations of the coarse-grain model, *τ_f,sim_*= 25sec, and previously experimentally measured *τ_f,exp_*= 28 sec (95% confidence interval: 26-31 sec, from the Fig. S13 in (Garzon-Coral et al., 2016)).

## DISCUSSION

We have shown that cortical pulling forces drive pronuclear migration and rotation, and spindle centering, elongation, and oscillations in *C. elegans* embryos, with no discernable contribution from MT pushing or cytoplasmic pulling forces. This was accomplished using a combination of FESLA, measurements of cytoplasmic fluid flow, detailed fluid mechanics simulations, and coarse-grained modeling. Thus, cortical pulling forces are sufficient to drive diverse cell biological behavior of centrosomes including directed motions, stable positioning, and oscillations. Since such centrosome behaviors are observed in diverse contexts (Cowan and Hyman, 2004, von Dassow et al., 2009, Longo and Anderson, 1968, Meaders and Burgess, 2020, Tanimoto et al., 2018, Garzon-Coral et al., 2016, Foe and von Dassow, 2008, Du and Macara, 2004, Riche et al., 2013, Zhu et al., 2013), as are cortical pulling forces (Wu et al., 2017), the principles we have uncovered here should be broadly applicable to other systems.

The coarse-grained model of centrosomes subject to cortical pulling forces that we have developed here is a constructive model, that is derived from the equations for dynamic instability of MTs (Dogterom and Leibler, 1993) with explicitly accounting for the geometry and mechanics of the interactions of MTs and cortical force-generators. Thus, this model allows for analytical predictions for how the behavior of centrosomes depends on the properties of MTs and cortical force generators. The coarse-grained model predicts that cortical pulling forces will stably position centrosomes when the rate of impingement of MTs onto cortical force generators is large enough: i.e. when the number and length of MTs emanating from the centrosome are sufficiently high. In this case, centrosomes will migrate from their initial locations until they reach the position at which they are stable – this provides an explanation for pronuclear centration in *C. elegans*. Once the centrosome reaches its resting position cortical pulling forces will actively maintain it at that position – this provides an explanation for the stable positioning of the spindle in *C. elegans*. Cortical pulling forces can stably position centrosomes because of two effects: a geometric effect in which pulling forces occur at increasingly oblique angles as the centrosome approaches the cortex; and, the stoichiometric interactions between MTs and force generators: i.e. each force generator can bind to at most one MT at a time (Farhadifar et al., 2020). If the time a MT stays attached to a cortical force generator is sufficiently small compared to the time it takes for a MT to grow across the cell, then the centrosome will start oscillating when the number of cortical force generator surpasses a critical value - this provides an explanation for anaphase spindle oscillations in *C. elegans*. The coarse-grained model provides analytical predictions for when these different regimes occur, as well as the speed of centrosome migration and the spring constant associated with spindle positioning.

Among the three proposed force generation mechanisms exerting on MTs, laser ablation of astral microtubules helps distinguish between pushing and pulling forces, while cytoplasmic fluid measurement differentiates between cortical pulling and cytoplasmic pulling. Our novel femtosecond stereotactic laser ablation (FESLA) enables us to accurately probe forces of specific orientations with minimal collateral damage. The motions that result immediately after ablation provide insight into the direction and origin of the perturbed forces.. The precise and highly controllable timing of ablation that FESLA enables studies of highly dynamic biological phenomena. In addition, our characterization of subcellular fluid flows was key to determining the nature of force generation. This was achievable because flows provide a signature of the mechanism of force generation. The detailed agreement between the flows that we experimentally observed and those predicted from our simulations shows that modern computational fluid dynamics methods can accurately describe cell biological systems. This validates the use of subcellular fluid flows as a broadly applicable method to study force origins. Overall, our integrated methodology of femtosecond stereotactic laser ablation (FESLA) and subcellular fluid flows are broadly applicable for studying intracellular and extracellular forces in biology.

## METHODS

### • LEAD CONTACT AND MATERIALS AVAILABILITY

Further information and requests for resources and reagents should be directed to, and will be fulfilled by the Lead Contact, Hai-Yin Wu.

### • EXPERIMENTAL MODEL AND SUBJECT DETAILS

#### ∘ Worm strains

The *Caenorhabditis elegans* line *AZ244 (unc-119(ed3) III; ruIs57[unc-119(+) pie-1::GFP::tubulin]*), and a new line obtained by crossing *AZ244* with a line expressing mCherry-labeled γ-tubulin, were used throughout this study. Additionally, to visualize chromosomes, *SA250 (tjIs54 [pie-1p::GFP::tbb-2 + pie-1p::2xmCherry::tbg-1 + unc-119(+)]; tjIs57 [pie-1p::mCherry::his-48 + unc-119(+)]*) were used for metaphase experiments shown in Figure 4A-4D. This line helped to determine the timing of metaphase and ensure that chromosomes were not ablated.

#### ∘ *C. elegans* culturing conditions and embryo preparation

Worms were grown on nematode growth medium (NGM) plates seeded with OP50 *Escherichia coli* bacteria, and incubated at 24°C. Gravid *C. elegans* hermaphrodites were dissected by needle tips to release their embryos into M9 buffer. Then mouth pipettes were used to pick and transfer early embryos onto flat 4% agarose (Bio-Rad) pads, which were prepared on microscopic slides (Walston and Hardin, 2010). The fresh made agarose pad was promptly trimmed to a size smaller than the area of a coverslip. After adding small amount of M9 buffer to keep embryos moisturized, the sample was covered with a coverslip and sealed swiftly for imaging. Under this mounting condition, embryos were slightly squeezed, and were held in place by the sunk agarose pad.

After embryo preparation were done, they were imaged either at 22°C or 18°C. For different laser ablation conditions, each set of embryos, including both control uncut and ablated embryos, were imaged at the same temperature for comparison (either all at 22°C or all at 18°C).

### • METHOD DETAILS

#### ∘ Experimental Methods

##### Microscopy

We imaged mounted C. elegans early embryos on an inverted microscope (Nikon, TE2000) using a 60X water-immersion objective (Nikon, CFI Plan Apo VC 60X WI, NA 1.2). Images were acquired using a spinning disk confocal unit (Yokugawa, CSU-X1) equipped with continuous-wave lasers for fluorescence excitation and an EM-CCD camera (Hamamatsu, ImagEM Enhanced C9100-13) for detection. GFP fluorescence was excited at 488 nm and collected through a bandpass filter with 514-nm center and 30-nm full width at half maximum (FWHM) wavelength. mCherry fluorescence was excited at 561 nm and collected through a bandpass filter with 593-nm center and 40-nm FWHM wavelength. Fluorescent nanodiamonds (FNDs) were excited at 561nm and collected through a long-pass filter with 647-nm cut-on wavelength.

Figure S1A indicates the terminology for embryo orientation and axis labels used throughout this manuscript: The x-y plane is defined as the imaging plane. The longitudinal direction (or the long axis) of the oblong embryo is assigned as the x-direction, while the transverse direction refers to the y-direction.

##### Femtosecond Stereotactic Laser Ablation (FESLA)

Our laser ablation setup was incorporated into an inverted microscope equipped with a spinning disk unit (Figure 1C). A near infrared-ray Ti:sapphire femtosecond pulsed laser beam with reduced repetition rate (16~80-KHz, compared to the common 80-MHz-repetition-rate output) (Chung et al., 2006, Chung and Mazur, 2009, Gabel et al., 2008, Gabel et al., 2007, Vogel et al., 2005) was directed and merged into the microscope light path through a dichroic mirror. The ablation laser pulses were focused onto the sample by the same high-numerical-aperture objective used for confocal imaging. We used two different methods to reduce the repetition rate of the femtosecond laser pulses: 1) Using a cavity-dumped Ti:sapphire laser (Cascade-1, KML, with 830~840-nm center wavelength and 20~30-fs pulse width) with either a 40-kHz pulse train with 2.5~4 nJ pulse energy (the energy measured under our 60X water-immersion objective), or an 80-kHz pulse train with 2.5~3.5 nJ pulse energy; 2) Using a pulse picker (Eclipse, KMLabs) to select a 16-kHz pulse train from the 80-MHz Ti:sapphire laser pulses (Mai-Tai, Spectra-Physics, with 800-nm center wavelength and ~70-fs pulse width).

We ablated complex patterns by moving the sample on a 3-axis piezo-stage (P-545 PInano XYZ, Physik Instrumente) and controlling laser exposure with a fast mechanical shutter (Newport). Ablation patterns and image acquisition were controlled with custom LabVIEW codes. The dimensions of FESLA ablated regions are displayed in supplementary figures: For experiments during late metaphase to anaphase spindle oscillations, see Figure S2A (for arc cuts in Figure 1E-1J), Figure S2E (for double-plane cuts), Figure S2F (for open cylinder cuts in Figure 2A-2E), and Figure S2I (for cup cuts in Figure 2F-2I); for experiments during metaphase, see Figure S4A-S4B (for plane cuts in Figure 4A-4B) and Figure S4C-S4D (for cup cuts in Figure 4C-4D); for experiments during pronuclear complex (PNC) stage, see Figure S4E (for plane cuts in Figure 5A-5B) and Figure S4H (for large semicircular arc cuts in Figure 5F-5H).

##### Fluorescent nanodiamonds (FNDs)

Passivated FNDs were used as tracer particles to visualize cytoplasmic flows. We adopted a 2-step surface treatment to coat FNDs with mono-methyl polyethylene glycol (mPEG). FNDs (40~50 nm in diameter, after acid wash treatment, a gift from Dr. Huan-Cheng Chang) (Fu et al., 2007, Mohan et al., 2010, Su et al., 2017, Vaijayanthimala et al., 2012) in Milli-Q water were sonicated for 30 mins, and then mixed with α-lactalbumin (Sigma, α-lactalbumin from bovine milk, calcium depleted) at a weight ratio of FND: α-lactalbumin = 1:2. The mixture was gently shaken overnight at 4°C, and then washed through a centrifugal filter (Amicon Ultra, 100 KDa) several times. The FND concentrate was dispersed in Milli-Q water again and sonicated in ice water bath for 15 minutes, and then optionally syringe filtered (Millex-VV, 0.1 μm, Durapore PVDF membrane). After diluting the solution of FND (with absorbed α-lactalbumin) to ≈ 0.1 mg/mL, boric acid was added to reach a final concentration of 5mM, and the pH was adjusted to 8. For the second mPEG coating, fresh mPEG-SVA (mPEG-Succinimidyl Valerate, M.W. 5000, Laysan Bio) was added to a weight ratio of FND: mPEG-SVA ≈ 1:1 and the mixture was stirred overnight at 4°C. Then the FNDs were washed through the centrifugal filter several times and dispersed in Milli-Q water. Immediately before use, the FND solution was sonicated in an ice water bath for 15 minutes, syringe filtered (Pall Acrodisc, 0.2 μm, HT Tuffryn membrane), and diluted to a concentration of 0.25~1 mg/mL.

We microinjected freshly coated FNDs (within 48 hours) into the distal gonads of *AZ244* (unc-119(ed3) III; ruIs57[unc-119(+) pie-1::GFP::tubulin]) young adult hermaphrodites using needles pulled from borosilicate glass capillaries (World Precision Instruments,1.0 mm OD, 0.58 mm ID, with filament) with a P-87 micropipette puller (Sutter Instrument). We cut the hermaphrodites 6~10 hours after microinjection, and retrieved their embryos for imaging.

We imaged *AZ244* embryos with injected FNDs at 18°C. Time-lapse z-stack confocal images of both GFP::β-tubulin and FND were acquired every 2.5~5 seconds. Centrosomes were visualized by GFP::β-tubulin images, which included several z-sections with 1-μm spacing and were taken around PNCs or spindles. FND signals were captured through z-stacks having 9 or 15 z-section images with 1-μm spacing.

#### ∘ Computational fluid dynamics simulations

##### I) Numerical methods for computing intracellular flows

We give a brief description of the computational methods used to compute the cytoplasmic flow results shown in the main text. These methods were developed and applied originally to study the dynamics of pronuclear migration and centering (Nazockdast et al., 2017b, Nazockdast et al., 2017c). These methods, based upon boundary integral methods, slender-body theory, and fast Stokes equation solvers, account for the hydrodynamic interactions that couple together microtubules (MTs), any internal bodies (here, the spindle or pronuclear complex (PNC), and centrosomes) and the cell cortex. They also account for MT flexibility, dynamic instability, and MT interactions with molecular motors or boundaries.

The cell interior is treated as having five structural elements (See Figure S5A): the spindle body or pronuclei, the centrosomes, the cell cortex (boundary), the arrays of MTs nucleated from the centrosomes, and the cytoplasmic fluid. Given the slow speed and small scale of intracellular flows, any inertial effects can safely be ignored. While we assume the cytoplasm is itself a Newtonian fluid (Garzon-Coral et al., 2016), the response of the MT-laden cytoplasmic system is non-Newtonian. The flow of a Newtonian cytoplasm is described by the incompressible Stokes equations:

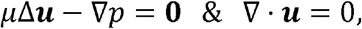

where *μ* is the cytoplasmic viscosity, ***u*** is the (cytoplasmic) fluid velocity, and *p* is the pressure which maintains flow incompressibility.

Solutions to the Stokes equations can be represented through boundary integral formulations (Pozrikidis, 1992), where the fluid velocity is represented as an integral distribution of fundamental solutions to the Stokes equations on all immersed and bounding surfaces. That is, for *N* MTs and *M* other immersed surfaces (cortex, centrosomes, …) we can represent these distributions in the form

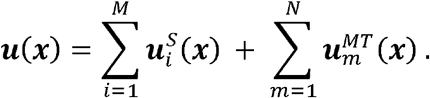

The densities of these distributions are determined and coupled together by the application of boundary conditions, such as the no-slip condition (surface velocity is equal to fluid velocity) or applied forces and torques. A boundary integral formulation reduces the computational problem from being three-dimensional (solving the Stokes equations in the fluid volume) to the two-dimensional problem of solving coupled singular integral equations on all the immersed and bounding surfaces.

Of particular interest here are MTs. MTs have a diameter of *a* ≈ 25 nm and lengths of *L* ~ 1 - 20 μm, yielding small slenderness ratios of *ϵ* = 10^−3^ - 10^−2^. This allows their flow contributions to be treated specially: their surface integrals can be reduced, through asymptotics, to line integrals of ‘’Stokeslet’’ fundamental solutions along their centerlines (Götz, 2001, Johnson, 1980, Keller and Rubinow, 1976). In particular, to leading order in slenderness ratio, the induced fluid velocity from the *m^th^* MT, having centerline coordinates ***X**_m_*(*s, t*), with *s* ∈ [0, *L_m_*] the arclength from its minus end and *L_m_* the MT length, is

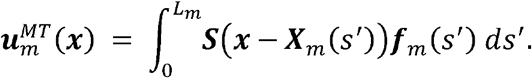

Here the second-rank tensor 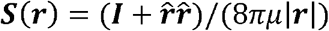, with 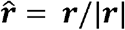, is the (single-layer) Stokeslet fundamental solution of the Stokes equations, and ***f_m_*** is the force/length the MT exerts upon the fluid. To leading order, the velocity of the *n^th^* MT centerline is given by

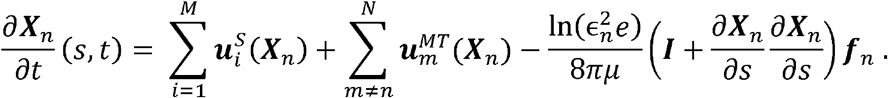

The force ***f**_m_* arises from internal elastic forces, and from external forces either at MT ends (through boundary conditions), or along MT lengths. The internal elastic forces are related to MT conformation by Euler-Bernoulli beam theory using the constitutive relation ***f*** = ‒*E**X**_SSSS_* + (*T**X**_S_*)*_S_*, with *E* the MT flexural modulus, and *T* is MT axial tension. The first term is the bending force per unit length and the second is the tensile force per unit length. The tension *T* is determined by the condition of MT inextensibility (Tornberg and Shelley, 2004).

MTs are nucleated from, and clamped rigidly to, centrosomes which are themselves mechanically coupled to the spindle or PNC. In our modeling, motion results from the application of forces and/or torques to these structural elements. The spatial discretization of all surfaces and MTs, application of quadrature formulae for surface integral equations, and the discretization in time of MT and body velocities results in a linear system of equations to be solved at each time-step to update all body positions and MT conformations. To solve this large system, we use the GMRES iterative solver (Saad and Schultz, 1986) with a block-diagonal preconditioner (Nazockdast et al., 2017c). Within this iterative method, we use a parallel implementation of the Kernel-Independent Fast Multipole Method (Malhotra and Biros, 2015) to hierarchically and efficiently evaluate all hydrodynamic interactions, leading to linear cost per time-step in the number of spatial unknowns (again, determined by the discretization of surfaces and MT center-lines).

###### I.i) Changes for the cytoplasmic pulling model

In the cytoplasmic pulling model, cargo-carrying dyneins walk along MTs towards the centrosomes (MTs’ minus ends) and so apply pulling forces on MTs (see Figure S5B). By Newton’s third law of motion, the pulling forces on MTs are equal in magnitude and opposite in direction to the force that dynein exerts on the cytoplasm to drag the cargo through it. We treat the density of the attached dynein as a continuum field with constant number of attachments per unit length of MT, and the motor force as aligned along the MT. Thus, the pulling forces on a MT per unit length is ***f**^motor^* = *F_dyn_n_dyn_**X**_s_* where *F_dyn_* is the magnitude of the force applied from a single motor to a MT, *n_dyn_* is the motor number density per unit length, and ***X**_s_* is the tangent vector to the MT. Accordingly, we modify the dynamics equation for MT motion to

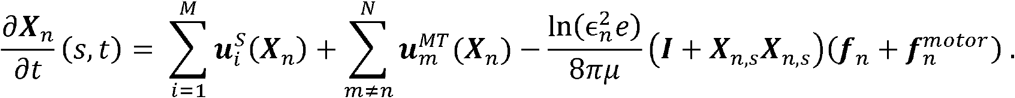

However, for induced flows away from the MT, the pulling forces on MTs are balanced by the forces exerted on the cytoplasm to drag the cargo (i.e. the far-field motor-induced flow is from a dipolar force), and the dominant effect is that from the internal elastic force ***f**_n_*. Thus, the expression for 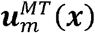 is unmodified (Nazockdast et al., 2017c, Stein et al., 2021).

##### II) Simulation setups

We used the experimentally measured centrosome velocities of the spindle assembly at various stages to set the conditions of comparison with our simulations. In particular, we performed short time simulations, explicitly to recover the instantaneous cytoplasmic velocity fields induced by motion of the spindle/pronuclei-centrosome assembly and the attached MTs.

In the cortical pulling model (Figure S5A), previous studies (Nazockdast et al., 2017b) show that MTs remain nearly straight under their extensile loading from cortically bound dynein motors. Hence, the cortical pulling forces directly act on the spindle/pronuclei-centrosome assembly without substantial loss to MT bending, which is equivalent to applying an external force and torque at the assembly’s center. In this case, the cytoplasmic velocity arises from the translations and rotations of the spindle/pronuclei-centrosome-MT complex (Nazockdast et al., 2017b). We implemented the cortical pulling mechanism by applying an external force and torque on the assembly which we adjusted to match the centrosome velocities in the experiments.

For the cytoplasmic pulling model, at short times as in our simulations, we find that the instantaneous velocities scale to a very good approximation with the magnitude of the applied pulling forces. Hence, we adjusted the motor force/length to match, in each case, the experimental centrosome velocities.

We also used experimentally measured positions and sizes of the spindle/pronuclei and centrosomes in the simulations (see Figure S5C). We modeled the centrosomes as rigid spheres of diameter 1 μm which, in most cases, were moved in a rigid frame with the pronucleus or spindle body. Each centrosome had on the order of 300 attached MTs. This number is smaller than the actual number of astral MTs (Redemann et al., 2017), however, by running the simulations for different numbers of MTs we found that the qualitative features remain similar.

Before starting the fluid dynamics simulations, we simulated MTs’ dynamic instability with a MT catastrophe rate of 0.015 sec^−1^, nucleation rate per centrosome of 62.5 sec^−1^ and MT growth velocity of 0.75 μm/sec. In particular, holding fixed the positions of the spindle/pronuclei and the centrosomes, we let MTs nucleate and grow from the centrosomes. Depolymerization occurred either through spontaneous catastrophe during free growth, or upon colliding with the cortex. After the MTs reached their steady-state length distribution (i.e., an exponential distribution truncated by catastrophe at the cortex), we activated either cortical or cytoplasmic pulling forces, depending upon the scenario. For each stage, we ran simulations for ten different initializations of MT arrays and presented averaged results over those samples.

###### II.i) Spindle in oscillation

At this stage, the spindle is modelled as a rigid ellipsoid of length 16.8 μm along the anterior-posterior axis (A-P axis), and of width 4 μm at the spindle center. The spindle was also slightly rotated in the counterclockwise direction (0.005 rad) because the anterior centrosome is below the A-P axis whereas the posterior one is upon it (see Figure 3E). The centrosomes are constrained to move in the same rigid frame as the spindle.

###### Cytoplasmic pulling

The cytoplasmic pulling model does not lend itself straightforwardly to capturing the observed centrosome and spindle speeds, and so adjustments are necessary. During spindle oscillation the spindle center is shifted towards the posterior of the cell. Hence, in our modeling of the centrosomal array, the posterior centrosome has shorter MTs towards the posterior end than does the anterior centrosome towards the anterior (see Figure S5C). Loading the MTs uniformly with motors then leads the spindle-centrosome assembly to move towards the anterior. Additionally, both centrosomes are almost on the A-P axis (see Figure 3E). Thus, the centrosomes have MTs of similar lengths above and below their centers, which generates weaker flow in the axis perpendicular to the A-P axis than that along the A-P axis. So, to match the centrosomes’ velocities, we varied the motor density across the MTs instead of having them uniformly distributed.

###### Cortical pulling

To capture the experimentally observed centrosome and spindle speeds, we applied forces in the x- and y-directions, and a torque in the z-direction, on the spindle-centrosome assembly.

###### II.ii) Spindle in prometaphase

We modeled the spindle in prometaphase as a sphere of 5 μm diameter. The experiment observations (see Figure 4G) show negligible motion of the centrosomes or spindle at this stage.

###### Cytoplasmic pulling

Since there is not any reference velocity for us to scale the motor force/length, we used the same value for all the MTs which is the same as in the pronuclear migration simulations. Their symmetry produces no motion of the centrosomes, but considerable cytoplasmic motion.

###### Cortical pulling

It is assumed that the total pulling force upon the spindle-assembly was zero, which generates neither centrosome nor cytoplasmic motion.

###### II.iii) Spindle in metaphase elongation

The spindle was modelled as a rigid ellipsoid of 12.5 μm in length, and of 4 μm in width at the spindle center. We modeled the elongation of the spindle as a prescribed rate of increase in distance between the centrosomes, as measured experimentally, with the spindle body constrained to stay at their midpoint.

###### Cytoplasmic pulling

Again, this model requires some adaptation to achieve the experimentally observed centrosome velocities. Here, the motor force/length was the same for all the MTs which has the same magnitude as in the prometaphase and pronuclear migration simulations. But, it was taken as slightly greater on the posterior MTs so as to prevent the anterior centrosome from moving. Due to greater motor force/length on the posterior MTs, one could expect to see faster streaming velocity on the posterior. However, this is not observed because the posterior centrosome’s motion in the x-direction drags fluid in the opposite direction to the flow the motor forces generate, and these counter-acting flows cancel each other near the posterior centrosome.

###### Cortical pulling

We let the centrosomes move as a result of the elongation of the spindle-centrosome complex along the A-P axis. Then, to reproduce the experimentally observed centrosome velocities, we applied a force on the spindle-centrosome assembly in the x-direction that canceled out the anterior centrosome’s velocity.

###### II.iv) Pronuclear migration

We modeled the pronuclei as a rigid sphere of 5 μm diameter. The centrosomes were constrained to move in the same rigid frame as the pronuclei.

###### Cytoplasmic pulling

We used the same motor force/length for all the MTs, keeping it similar in magnitude to those used in the other simulations, but adjusting it to achieve the observed centrosome velocities.

###### Cortical pulling

To achieve the observed centrosome velocities, we applied a torque in the clockwise direction and a force with components in the anterior and y-directions.

###### II. v) Combining the pulling models

In the both cortical and cytoplasmic pulling models (Figure S5), previous studies (Nazockdast et al., 2017b) show that MTs remain nearly straight under their extensile loading by dynein motors. Hence, in our computations here, again, we take the MTs to be straight for both models and so contributions from bending forces are absent. The consequent velocity field will then be that for a rigid structure of MTs, centrosomes, and PNC or spindle. That velocity field is then found through solution of linear equations for MT tension *T*, and the surface vector density **q** of the double-layer integrals that captures the velocity contributions from the immersed surfaces of the centrosomes, PNC/spindle, and cortex (Nazockdast et al., 2017b).

For the cortical pulling model these equations can be written in the form

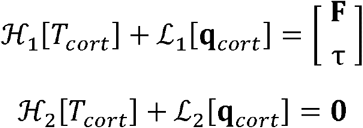

where **F** and **τ** are the applied force and torque, and 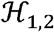 and 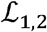 are linear integral and differential operators acting along MTs and all surfaces. For the cytoplasmic fulling model the equations instead have the form

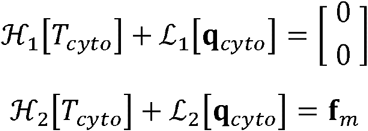

where **f***_m_* is comprised of the vectors *f_m_***X***_m_* of motor forcing. Note that in either case that if **F = τ = 0**, or **f***_m_* = 0, then the tension and density will both be zero.

Given a tension *T*, and a density **q**, the cytoplasmic velocity can generally be expressed as

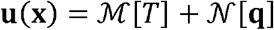

Where 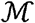 and 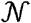 are again linear integral and differential operators. Now, if instead we consider a mixed model, with contributions from both cortical and cytoplasmic pulling, of the form

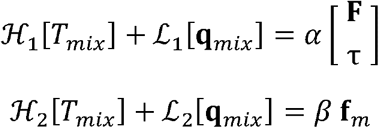

then *T_mix_* = *αT_cort_* + *βT_cyto_* and **q***_mix_* = *α***q***_cort_* + *β***q***_cyto_* is its solution. Hence, the induced cytoplasmic velocity, using linearity, is

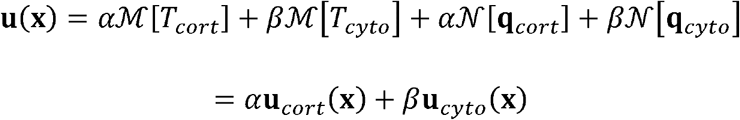

that is, is a linear combination of both cortical and cytoplasmic pulling models. As a verification on our code, we have directly checked this property of a mixed pulling model.

#### ∘ Coarse-grained model of cortical pulling

Here we derive a coarse-grained model of a centrosome moving in a spherical cell, subject to pulling forces from force generators on the cell surface. We start by considering an isolated centrosome nucleating microtubules (MTs) that undergo dynamic instability (*section I*). We then construct a model of stoichiometric force generators in which each force generator can bind to at most one MT at a time (*section II*). We show that pulling forces from stoichiometric force generators are sufficient to stably position the centrosome or produce oscillations (even in the absence of additional forces from MT pushing or bending). We next construct a model of non-stoichiometric force generators in which force generators bind all MTs that contact them (*section III*). We show that pulling forces from non-stoichiometric force generators are always destabilizing and cannot stably position the centrosome.

##### I) Nucleation and growth of microtubules (MTs) from a centrosome

We consider a centrosome with MTs nucleating with equal probability in all directions with rate *γ*, which then grow with velocity *V_g_* and undergo catastrophe with rate *λ*. The length distribution of MTs, *ψ*(*l, t*), satisfies the Fokker-Planck equation:

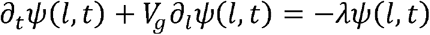

Setting the flux *V_g_ψ* at *l* = 0 to be equal to the nucleation rate gives *V_g_ψ*(0, *t*) = *γ*, or *ψ*(0, *t*) = *γ/V_g_*. Starting from an initial state with no MTs present (i.e. *ψ*(*l* > 0,0) = 0), the complete time-dependent solution for the length distribution of MTs is

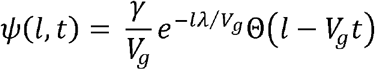

where Θ(*a*) = 1 for *a* ≤ 0 and Θ(*a*) = 0 for *a* > 0. Thus, there is a front of MTs propagating outwards at a speed *V_g_*. No MTs extend beyond the front, while MTs behind the front have a (truncated) exponential length distribution. At long times, this approaches the steady state MT length distribution 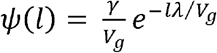.

The total number of MTs, *N_MT_*(*t*), is given by 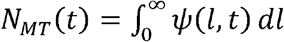, which, for an unconfined centrosome at steady-state, becomes 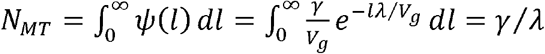.

##### II) Stoichiometric force generators

###### II.i) An individual stoichiometric force generator

Here we derive a model for stoichiometric force generators, in which each force generator can bind to at most one MT at a time. An unbound force generator will bind to a MT that contacts it, while a bound force generator detaches from the MT it is associated with at rate *κ*. When the force generator is bound to a MT, it exerts a pulling force on the centrosome of 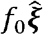, where 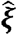 is the unit vector pointing from the centrosome to the force-generator (Figure 6B, left). Thus, the expected force that the force generator exerts on the centrosome at time *t* is 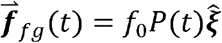, where *P*(*t*) is the probability that the force generator is bound to a MT at time *t*. *P*(*t*), in turn, obeys the dynamics

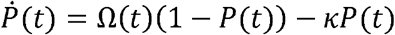

Where Ω(*t*) is the rate at which MTs impinge upon the force generator at time *t*. We next calculate Ω(*t*).

###### MT impingement rate

We calculate the rate that growing MTs impinge upon a disk-shaped force-generator of radius *r*, located a distance *d* away from a centrosome at the origin. Only MTs located in a cone defined by the position of the centrosome and the projected area of the force-generator can grow to contact the force-generator. The number of MTs located in this cone is (approximately)

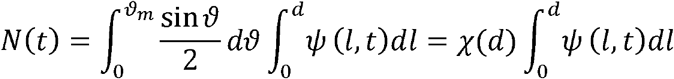

where *ϑ_m_* = tan^−1^ (*r/d*) is the solid angle of the cone and 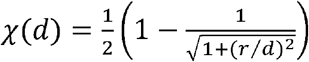 is the fraction of MTs nucleated from the centrosome that fall inside the cone.

For the moment we shall assume that growing MTs are impinging directly upon the force generator, i.e. that *ψ*(*d, t*) > 0, and that the centrosome is moving at speeds slower than *V_g_*. The time derivative of the number of MTs inside the cone is then

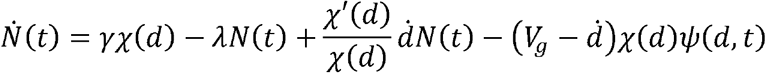

The first term is gain from nucleation and the second term is loss from catastrophes. The third term is change from the cone getting wider or narrower from changes in *d*. The last term is a loss term from MTs hitting the force generator at the cone end, a distance *d* from the centrosome. If the centrosome is moving with speed 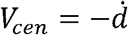, either directly away from or to the force generator, then the rate of impingement of MTs is given by

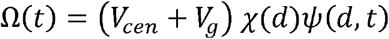

and since, by assumption, |*V_cen_*| < *V_g_*, this rate is always positive. More generally, if the centrosome is moving with velocity 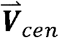, then the plus ends of MTs in the cone move with net velocity 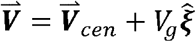, and the rate of impingement of MTs on the force generator is

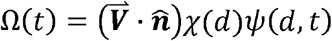

where 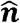 is an outward unit vector normal to the surface of the force generator. At any time *t* > *d/V_g_*, after the onset of nucleation, one can assume that 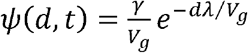.

In this model, MTs grow until they hit the cell boundary at which point they either bind to an unoccupied force generator, or disassemble. If the centrosome moves away from the force generator at speeds greater than *V_g_* (i.e. 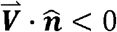), then the growing MT plus ends will have a net velocity that also causes them to move away from the force generator, and hence the rate of impingement Ω(*t*) of these MTs onto the force generator will be zero. As described in *section I*, these MTs will form a free front of plus-ends growing away from the centrosome at speed *V_g_*. The evolution of that front, and determining when it reattaches to the boundary, becomes part of the full dynamics and is discussed in *section II.v* on nonlinear dynamics.

###### II.ii) Net pulling force in a spherical cell with stoichiometric force generators

We next calculate the net pulling force acting on a centrosome in a spherical cell of radius *W* with *M* stoichiometric force generators uniformly spread over the cell surface (Figure 6B, right). We consider a centrosome moving along the y-axis, located at position *Y*(*t*), moving with velocity 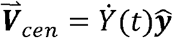. Again, for the moment we assume that the centrosome is moving at speeds slower than *V_g_*.

The net force acting on the centrosome is obtained by summing the force from all *M* force generators, each located at position 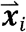

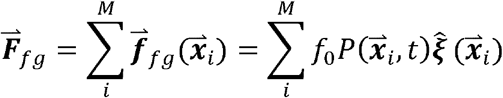

To make further progress, we coarse-grain by approximating the sum over discrete force generator positions by an integral over all positions on the surface of the sphere (which is equivalent to assuming a continuum of force generators, uniformly covering the sphere)

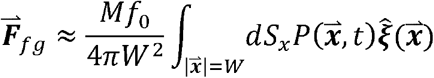

We assume symmetry about the y-axis and parameterize the location of the force generators by the polar angle *φ* (Figure 6B). A force generator at angle *φ* on the surface of the cell is located at position 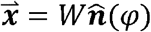, where 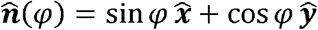 is the unit vector normal to the surface of that force generator. The vector from the centrosome to the force generator is 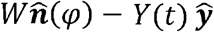, so the unit vector pointing from the centrosome to the force-generator is 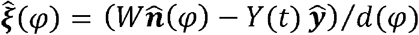. The distance from the centrosome to the force generator is 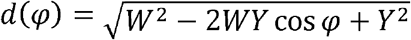. Thus, the MT plus ends move at velocity 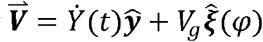, and the projection of the MT plus end velocity onto the direction of the force generator is 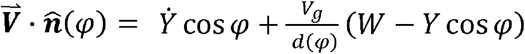.

Then, projecting the net pulling force in the 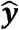 direction gives

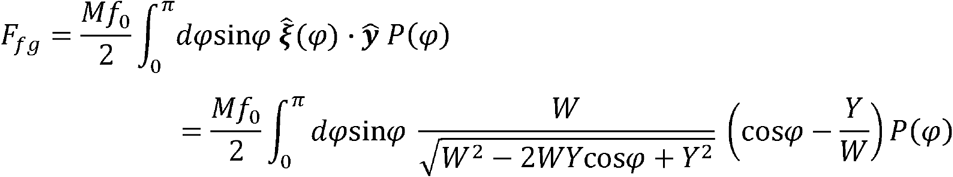

###### II.iii) Equations of motion with stoichiometric force generators

In addition to pulling forces from the force-generators, we also consider the drag on the centrosomes, 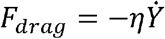, with drag coefficient *η*. From force-balance, *F_drag_* + *F_fg_* = 0, which gives

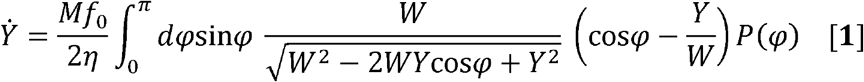

The equations of motion for the system consist of two coupled equations, the force-balance equation [**1**] for 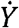, and the dynamical equation for *P*:

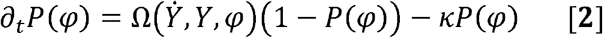

where

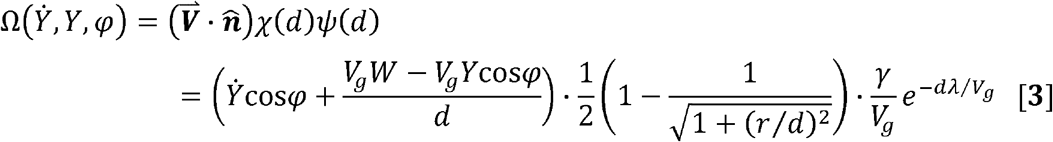

is the impingement rate, and

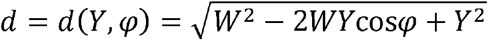

###### II.iv) Centered steady-state and its stability with stoichiometric force generators

At steady-state, 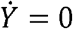 and 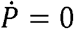, which results when the centrosome is at the center of the sphere, with steady-state position 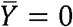, and steady-state attachment probability

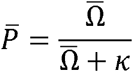

Where the steady-state rate of impingement of MTs onto the surface of the sphere is

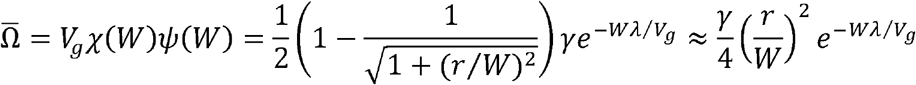

We next investigate the stability of the centered state by considering small perturbations: i.e. 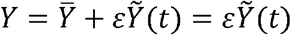 and 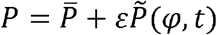 for *ε* << 1. Inserting the perturbations into the equations for 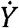 and 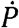, retaining only the leading-order *O*(*ε*) terms, and integrating the 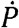 equation against sin*φ* cos*φ*, yields two ODEs:

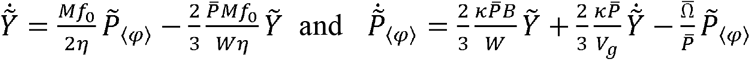

Where 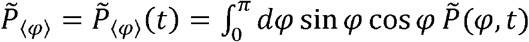 and 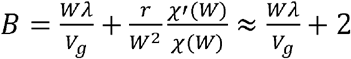. The approximate expression for *B* omits terms of order 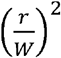, which can be safely neglected because the force-generators are significantly smaller than the cell in all cases of interest. Taking a time derivative of the 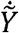 equation, and substituting back in the 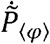 equation, leads to a single second-order ODE for this system:

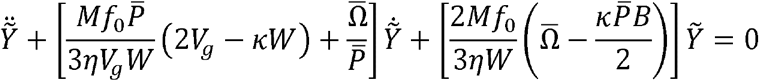

We search for solutions of the form 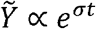, yielding

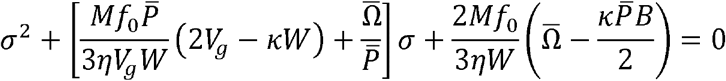

This gives two roots. If the real parts of both roots are negative, then the centrosome will stably position in the center of the cell. Starting from stable centering, we consider two ways to lose stability: 1) if both roots remain real, but one becomes positive, then the centrosome will tend to spontaneously decenter; 2) if both roots become purely imaginary, then the centrosome undergoes a Hopf bifurcation to an oscillatory state. We next perform a more detailed examination of three cases −spontaneous decentering, oscillations, and stable centering.

###### Spontaneous decentering

If 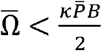 then both roots are real, and at least one of them is positive. Under this scenario, the centrosome will tend to spontaneously decenter. This criterion can be roughly interpreted as stating that the center will lose stability if the rate that unbound motors become bound, 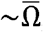, becomes less than the rate that bound motors unbind MTs, 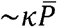. The inequality can be approximated as 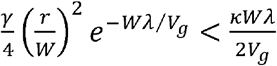. Thus, the center becomes unstable when the nucleation rate, *γ*, size of the force generators, *r*, or average MT length, *V_g_/λ*, becomes too small; when the size of the cell, *W*, increases too much; or when the detachment rate, *κ*, becomes too large. Note, that in the case considered here, of non-interacting force generators, the criterion for spontaneous decentering is independent of *M*, the number of force generators. Thus, the centrosome can remain stably positioned with arbitrarily more force generators than MTs, demonstrating that the stability of the center is not due to a “limiting number” of force generators.

###### Oscillations

When 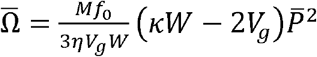 the eigenvalues are purely imaginary, i.e. *σ* = ±*σ_c_* = ±*iω_c_*, which indicates oscillatory behavior. Passing from the stable centering regime, described above, through this point to an oscillatory state is a loss of stability via a Hopf bifurcation. This Hopf bifurcation can only occurs if 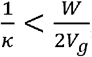, i.e. only if the time a MT stays attached to a force generator is sufficiently small compared to the time it takes for a MT to grow across the cell. In that case, increasing motor number causes such a transition from stable centering to oscillations at a critical number of motors

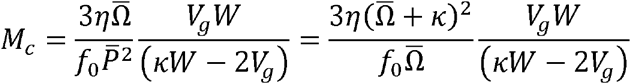

with a frequency

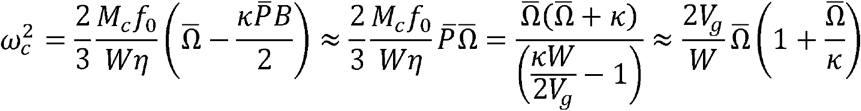

###### Stable centering

The centrosome is stable in the center position if the real parts of both roots are negative. This occurs when the rate at which MT impinge upon the cell surface is sufficiently high. Specifically, it requires that

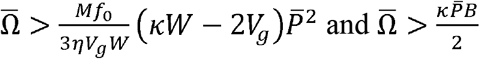

When the centrosome is stably positioned at the cell center, it is natural to ask how it responds to an externally applied force. Force-balance then becomes *F_drag_* + *F_fg_* + *F_ext_* = 0. Taking the external force to be small, *F_ext_* = *εf_ext_*, and expanding to linear order yields

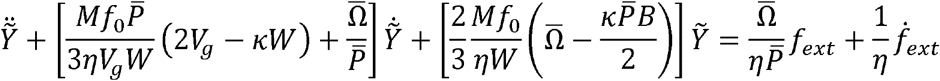

Note that the response of the centrosome to an applied force is not equivalent to a spring-and-dashpot model because the additional degree of freedom associated with motor attachments makes the dynamics of 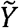 contain a second derivative with respect to time and explicitly dependent on the rate of change of the applied force (i.e. it is a second order system with numerator dynamics). However, at steady-state, this becomes

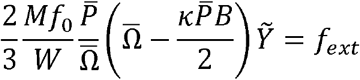

Thus, at steady-state, the displacement, 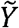 varies linearly with the applied force, like a spring, with spring-constant

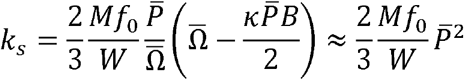

If the center position is stable then, 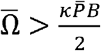, and thus *k_s_* > 0, as expected. Note that the spring constant is roughly the average number of engaged force generators, 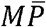, times the average force per force generator, 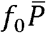, divided by the width of the cell, *W*.

If the centrosome is displaced from the center (or if it starts off center), then it will move to the cell center. After a quick initial transient, the approach to the center will be exponential, with a time scale, *τ_c_*, given by

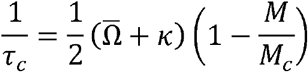

This same time-scale dominates in the response to an applied force. Note that a naïve guess for *τ_c_* would be the spring constant, *k_s_*, divided by the drag coefficient, *η*, as would result from a simple spring and dashpot mechanical model. However, in our model, based on the biophysics of MTs and molecular motors, *τ_c_* is not simply related to the spring constant, but rather depends on processes such as MT detachment rate and polymerization dynamics.

###### II.v) Non-linear dynamics with stoichiometric force generators

We next consider the full nonlinear, time-dependent behavior of a centrosome in a spherical cell with stoichiometric cortical force generators (including allowing for centrosome speeds greater than *V_g_*). As described in *section I*, the full time-dependent solution of *ψ*(*l, t*) contains a front of MTs propagating outwards at a speed *V_g_*. If this front of leading MT plus-ends is not in contact with a force generator, then Ω(*t*), the impingement rate of MTs upon that force generator, is zero. If the front does contact the force generator, then the impingement rate is given by the value calculated in *section II.i*. With this procedure for determining Ω(*t*), we numerically solve the equations of motion for 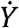 and 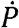 given in *section II.iii*, while numerically tracking the front of leading MT plus-ends.

We describe the front of leading MT plus-ends in terms of a coordinate system centered on the centrosome with polar angle *φ*′. There is a one-to-one mapping between this coordinate system and the coordinate system used to describe the position of the motors, which is centered on the cell center with polar angle *φ*. Thus, we can write *φ* = *φ*(*φ*′, *t*). We assume that the centrosomal array does not rotate, and represent the location of the plus-end front as 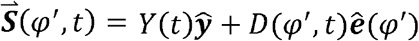, where 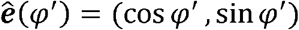 and *D*(*φ*′, *t*) is the distance from the centrosome to the front. The front is in one of two states; either

i. the front is at the cortex, i.e. *D*(*φ*′, *t*) = *d*(*φ*(*φ*′, *t*), *t*). So, MTs are impinging on force generators, which implies Ω(*φ*(*φ*′, *t*), *t*) > 0; or
ii. the front is short of, and growing towards, the cortex, i.e. *D*(*φ*′, *t*) < *d*(*φ*(*φ*′, *t*), *t*) and *∂_t_D*(*φ*′, *t*) = *V_g_*. So, MTs are not impinging on force generators, which implies Ω(*φ*(*φ*′, *t*), *t*) = 0.

Moreover, the front remains in contact with the cortex, i.e. in state (i), only so long as the centrosome is not moving away faster than *V_g_*.

We numerically track the front through a first-order time-stepping framework: Initial data *Y*^0^, *P*^0^(*φ*), and *D*^0^(*φ*′) are specified. Then,

1. given a time-step Δ*t*, and (*Y^n^, P^n^, D^n^, Ω^n^*) (and hence *d^n^*) at time *t^n^* = *n*Δ*t* find *Y*^*n*+1^ and *P*^*n*+1^ at time *t*^*n*+1^ = (*n* + 1)Δ*t* using Euler’s method for the equations of motion for 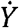 and 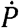 given in *section II.iii*.
2. Let 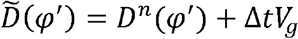; Given *Y*^*n*+1^ determine the mappings *φ* = *φ*(*φ′, t*^*n*+1^) and its inverse *φ*′ = *φ*′(*φ*, *t*^*n*+1^), and *d^*n*+1^(*φ**).
3. Update *D* and Ω according to the following scheme:

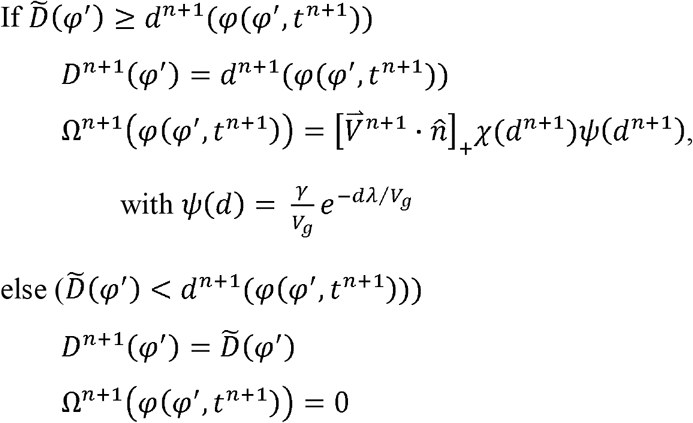

this completes one time-step.

##### III) Non-stoichiometric force generators

###### III.i) An individual non-stoichiometric force generator

Here we derive a model for non-stoichiometric force generators, in which any MT that contacts a force-generator is subject to pulling forces, even if that force-generator is already pulling on other MTs. A MT that contacts a force generator binds to it, while bound MTs detach from force generators at rate *κ*. Every MT bound to the force generator is subject to a pulling force 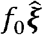, where 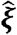 is the unit vector pointing from the centrosome to the force-generator (Figure 6B). Thus, the force that the force generator exerts on the centrosome at time *t* is 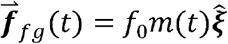, where *m*(*t*) is the number of number of MTs bound to the force generator at time *t*. The expected value of *m*(*t*), in turn, obeys the dynamics

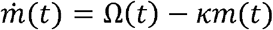

where Ω(*t*) is the rate at which MTs impinge upon the force generator at time *t*. As derived in *section II.ii*,

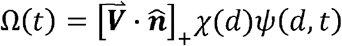

Here [*a*]_+_ = *a* for *a* > 0 and zero otherwise, and its appearance here reflects the fact that MTs will not reach the force generator if the centrosome is moving away faster than the MT growth speed. 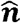 is a unit vector normal to the surface of the force generator, *d* is the distance between the force generator and the centrosome, and the net velocity of the plus ends of MTs is 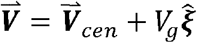 (where 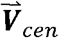 is the velocity of the centrosome).

###### III.ii) Net pulling force in a spherical cell with non-stoichiometric force generators

We next calculate the net pulling force acting on a centrosome in a spherical cell of radius *W* with *M* non-stoichiometric force generators uniformly spread over the cell surface (Figure 6B). We consider a centrosome moving along the y-axis, located at position *Y* (*t*), moving with velocity 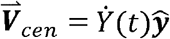. The net force acting on the centrosome is obtained by summing the force from all *M* force generators, each located at position 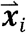

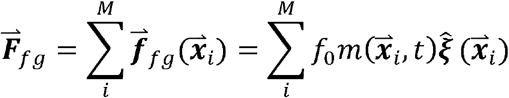

To make further progress, we coarse-grain by approximating the sum over discrete force generator positions by an integral over all positions on the surface of the sphere (which is equivalent to assuming a continuum of force generators, uniformly covering the sphere)

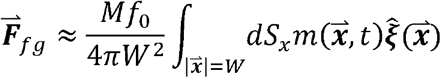

Projecting this net pulling force in the 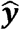 direction, and using the geometry described in *section II.ii* gives

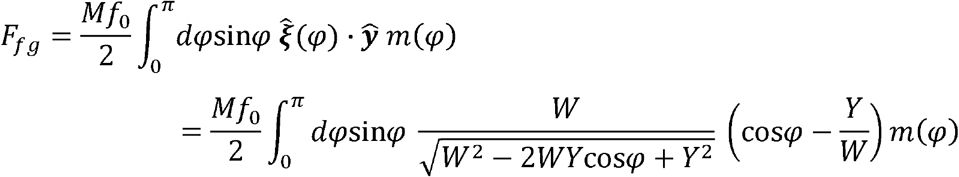

###### III.iii) Equations of motion with non-stoichiometric force generators

In addition to pulling forces from the force-generators, we also consider the drag on the centrosomes, 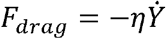, with drag coefficient *η*. From force-balance, *F_drag_* + *F_fg_* = 0, which gives

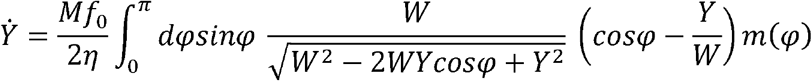

The equations of motion of the system consist of three coupled equations, this force-balance equation for 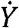, the dynamical equation for 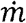, and the evolution equation for the MT length distribution, *ψ*:

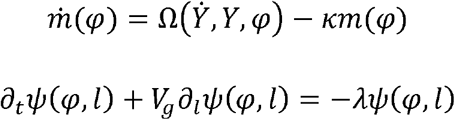

where the MT length distribution, *ψ*, can vary with angle around the centrosome. *ψ*(0, *t*) = *γ/V_g_* and *ψ* is coupled to the boundary by the condition that MTs do not grow past the cell surface. *m* is coupled to *ψ* through the rate of impingement of MTs

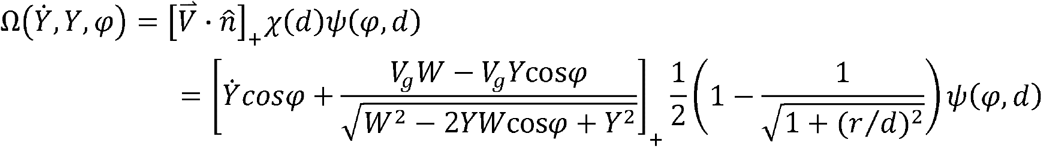

and

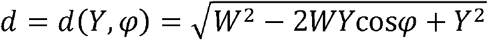

###### III.iv) Steady-state and stability with non-stoichiometric force generators

At steady-state, 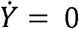 and 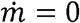, which results when the centrosome is at the center of the sphere, with steady-state position 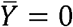, and steady-state number of attached MTs per force generator 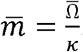. Where the steady-state rate of impingement of MTs onto the surface of the sphere is

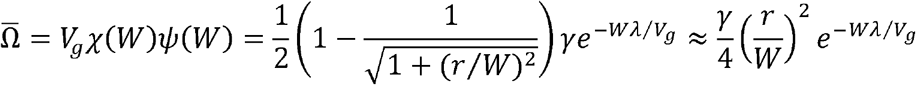

We next investigate the stability of the centered state by considering small perturbations: i.e. 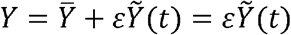 and 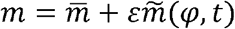 for *ε* << 1. Inserting the perturbations into the equations for 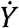 and 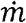, retaining only the leading-order *O*(*ε*) terms, and integrating the 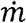 equation against sin*φ* cos*φ*, yields two ODEs:

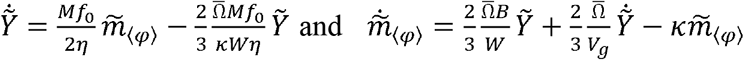

Where 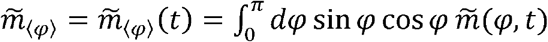 and 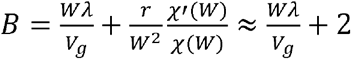. The approximate expression for *B* omits terms of order 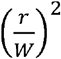, which can be safely neglected because the force-generators are significantly smaller than the cell in all cases of interest. Taking a time derivative of the 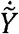 equation, and substituting back in the 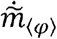 equation, leads to a single second-order ODE for this system:

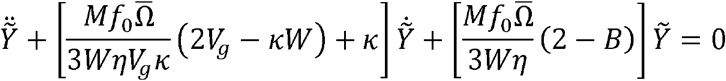

We search for solutions of the form 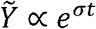, yielding

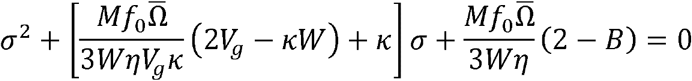

This gives two roots. Note that since *B* > 2, one of the roots is always positive: i.e. small perturbations will grow and the center position is generically unstable. Thus, non-stoichiometric force generators produce pulling forces that are destabilizing. In contrast, pulling forces from stoichiometric force generators can result in stable centering (see *section II.iv*).

#### • QUANTIFICATION AND STATISTICAL ANALYSIS

All image analyses and data quantification for experimental data were done in MATLAB (MathWorks).

##### ∘ Tracking centrosomes

The locations of centrosomes were extracted from either fluorescent *γ*-tubulin or fluorescent β-tubulin (Figure S1B). For *γ*-tubulin images (Figure S1B upper right), we adopted the particle-finding codes from MATLAB Particle Tracking Code Repository by Daniel Blair and Eric Dufresne (http://site.physics.georgetown.edu/matlab/) (adapted from IDL Particle Tracking software from David D. Grier, John C. Crocker, and Eric R. Weeks) (Crocker and Grier, 1996) to find the centrosome locations. We used high-pass background subtraction followed by thresholding and identification of local maxima. Sub-pixel localization of the anterior and posterior centrosome (orange and blue circles on Figure S1B upper right image) was achieved by calculating the centroid near the identified local maximum. For β-tubulin images, the tracking algorithm described above worked well at the pronuclear complex (PNC) stage, but the complex appearance of spindle centrosomes (Figure S1B lower right image) necessitated an alternative approach. In this case, we used an image correlation based approach: we constructed a square region with an artificial ring with the size similar to a centrosome (Figure S1C, upper), convolved this “synthetic” centrosome against background-subtracted and thresholded images of spindles (Figure S1C, lower), and identified the position of centrosomes as local maximum of this correlation (labeled circles on Figure S1B lower right image). After acquiring the locations of the centrosomes in each frame (for either *γ*-tubulin or β-tubulin imaging), we then used the tracking routines from the MATLAB Particle Tracking Code Repository by Daniel Blair and Eric Dufresne (http://site.physics.georgetown.edu/matlab/) (adapted from IDL Particle Tracking software from David D. Grier, John C. Crocker, and Eric R. Weeks) (Crocker and Grier, 1996) to link these positions into trajectories.

During centrosome oscillations, we defined the longitudinal direction of motion to be along the best fit straight line to the combined trajectories of the two centrosomes (Figure S1D). During metaphase and PNC stage, and for certain ablation data during oscillation, we instead defined the longitudinal direction of motion by the embryo’s geometry: we determined the cell boundary by thresholding (and smoothing) the β-tubulin image (Figure S1E, purple), fit the contour to an ellipse (Figure S1E, dashed yellow), and used the long axis of the ellipse as the longitudinal direction (Figure S1E, solid yellow line). We defined the location of the anterior and posterior ends of the embryo to be the location on the embryo contour that intersected with this longitudinal axis (Figure S1F). We defined the transverse direction to be orthogonal to the longitudinal direction.

##### ∘ Analysis of spindle oscillations

###### Centrosome motions

To determine the amplitude and timing of centrosome oscillations, we examined the transverse position of the centrosomes, obtained rough peak positions by identifying local maximum and minima after smoothing the trajectories, and then obtained refined peak positions by fitting the unsmoothed transverse position raw data around these local maxima/minima to Gaussians (Figure S1G).

We analyzed the impact of laser ablation arc cuts (Figure 1E-1J and Figure S2A) on oscillations by calculating *Δv_y_* = *v_y(after)_* - *v_y(before)_*, the difference in centrosome (y-)velocity immediately before, *v_y(before)_*, and after ablation, *v_y(after)_* (Figure S2B). Ablation itself took a time window, *τ_w_*, of 2.3 seconds on average to complete. Velocities were measured by linear fits of transverse centrosome positions over for a mean of 2.7 seconds (at least 3 data points, Figure S2C for example). The resulting centrosome transverse velocity changes (*Δv_y_* = *v_y(after)_* - *v_y(before)_*) were calculated for all arc-cut experiments shown in Figure 1E, 1F, 1H, and 1I (*Δv_y_* shown in Figure S2D).

We analyzed the impact of laser ablation double-plane cuts (Figure S2E) and open cylinder cuts (Figure 2A-2C and Figure S2F) on oscillations by calculating *A_after_/A_before_*, the ratio of amplitudes for the oscillations immediately before, *A_before_*, and after, *A_after_*, the cuts (Figure S2G). Both double-plane cuts and open cylinder cuts exhibited highly significant difference in the *A_after_/A_before_* ratio from controls (Figure S2H).

We analyzed the impact of laser ablation cup cuts (Figure S2I) on oscillations by calculating the minimum distance *d_min_* that the centrosome came to the cell boundary after ablation (Figure S2J).

###### Analysis of astral microtubules (MTs)

To measure the changes in astral MTs during spindle oscillations, we imaged GFP::β-tubulin in 11 embryos and calculate the intensity of pixels in three annular sectors with 3-μm inner radii, 9-μm outer radii, and 60° subtended angles (Figure S7A): an “up” transverse section oriented in the direction of centrosome motion, a “down” transverse section oriented in the direction opposite centrosome motion, and a “lateral” section between the centrosome and the lateral cortex. The orientations of the annular sectors remain unchanged while their locations vary as the posterior centrosome moves (Movie S7). For each embryo, we measured the tubulin intensity with background subtraction in these three regions throughout the 5-half-cycle temporal window with the maximal 5 consecutive peak-to-valley amplitudes (see Figure S6B left panel for example). Two full cycles of the upward part (positive y-direction, *T_1_* and *T_2_*), and two full cycles of the downward part (negative y-direction, *T_3_* and *T_4_*) after vertical flipping were taken into consideration (Figure S6B middle panel), giving a total of 44 full oscillation cycles from 11 embryos. For each full cycle, the time period was rescaled to the mean time period *T_(m)_* of the 44 cycles (Figure S6B right panel), and the intensity values from the transverse region were further normalized by the temporal mean intensity of the lateral region. After aligning the oscillation midpoints at zero, all amplitude values were divided by half the mean transverse displacement (maximum y-value minus minimum y-value) of the 44 cycles, to set the amplitude nearly between 1 and −1.

To analyze the predicted changes in astral MTs during spindle oscillations from the coarse-grained model, we derived an analytical formula for the total length of MTs in an annular sector (which is analogous to the experimentally measured intensity of tubulin in such a sector). To do this, we consider an angle *θ*′ ∈ [-*π, π]* encircling the centrosome in the coarse-grained model, where *θ*′ relates to the polar angle *φ*′ (defined in *section II.v* of the Coarse-grained model of cortical pulling portion of the Star Methods) by *θ’ = φ*′ for *θ*′ > 0 and *θ’ = −φ*′ for *θ*′ < 0. Consider a diskal sector 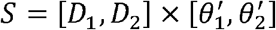 where we assume *D*_1,2_ < *d*(*θ*’) for 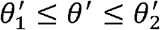. There are three types of MTs emanating from the centrosome: (1) those with length *l* ≤ *D*_1_ which contribute nothing to the diskal intensity; (2) those with length *D*_1_ ≤ *l* ≤ *D*_2_ which contribute a length of *l* – *D*_1_, and (3), those with length *D*_2_ ≤ *l* ≤ *D*(*θ*′) which contribute a length of *D*_2_ – *D*_1_. Hence, the total MT length in *S* is

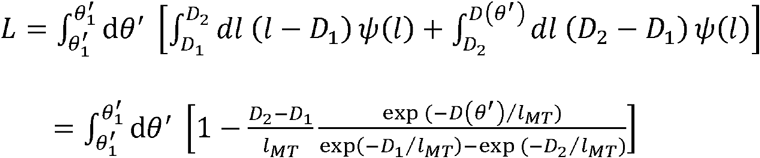

with *l_MT_* = *V_g_/λ*, and using that 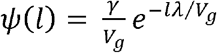.

##### ∘ Measuring cytoplasmic fluid flows with fluorescent nanodiamonds (FNDs)

We measured cytoplasmic fluid flows by using FNDs as passive tracer particles. This entailed: 1) tracking the motions of individual FNDs; 2) averaging the velocities of FNDs over multiple time points and embryos to determine fluid flow velocities.

We imaged FNDs in embryos by 3D time-lapse microscopy with either 9 or 15 z-sections spaced 1-μm apart. We adopted and modified the MATLAB 3D feature finding algorithms, which were written by Yongxiang Gao and Maria Kilfoil (Gao and Kilfoi, 2009) based on IDL codes from John C. Crocker and David G. Grier (Crocker and Grier, 1996), to extract the FND particle features and obtain their 3D positions with sub-pixel precision. Then we tracked the 3D trajectory of individual FND again using MATLAB Particle Tracking Code Repository by Daniel Blair and Eric Dufresne. We calculated the 3D instantaneous velocity of every FND, at each time and location, as the difference between its positions at two consecutive frames divided by the time interval between those frames. While tracking FNDs, we also imaged and tracked centrosomes using GFP::β-tubulin images.

To determine the pattern of fluid flow throughout the cytoplasm, we averaged together FND velocities from different time points and embryos. Positioning stages can mainly be recognized through GFP::β-tubulin images. We also used the position and movement of centrosomes to determine the frames to include and the coordinate system with which to perform this averaging. For spindle oscillations, we focused on the time, T_m_, where the centrosome is moving fast through the midpoint of its oscillation (Figure 3E). We selected times in which the posterior centrosome was located within 1 μm transversely from the oscillation midpoint, moving with a transverse speed greater than 0.15 μm/sec. For prometaphase spindles, we included times after rotation had nearly ceased. For metaphase spindles, we included spindles with lengths between 20% ~ 40% of their embryo lengths and before spindle oscillations began. After determining which time points to include, we aligned position and velocity data based on the locations of the centrosomes and the embryos’ anterior-to-posterior directions. Then we projected the 3D positions and instantaneous velocity vectors of the FNDs onto the x-y plane, and determined the 2D fluid flow velocity on grid points with 2.5-μm(x) × 2.5-μm(y) spacing (Figure S3A) by averaging FND velocities in overlapping cuboid domains. From prometaphase to late anaphase (Figure 3E, 6G, and 6J), the averaging domain was a 5-μm(x) by 5-μm(y) by 6-μm(z) cuboid with the grid point at the cuboid center (Figure S3B). To calculate fluid flows around pronuclei (Figure 5E), we used a 5 μm(x) by 5 μm(y) by 9 μm(z) overlapping averaging domain (Figure S3C). For PNC migration and rotation, we considered two different criteria for averaging: 1) As in the main text (Figure 5E), we adopted times when the longitudinal distance between the PNC center and the embryo posterior end measured 35% ~ 50% of the embryo length (or *R* = 0.35~0.5) (Figure S3D, lower left); 2) alternatively, we used times when the angle *A*, between the PNC axis (centrosome-to-centrosome) and the longitudinal axis of the cell, ranged between 33° and 66° (Figure S3D, lower right). Both methods gave qualitatively similar results.

The averaged experimental velocity vectors were plotted according to a *p*-value-related color map. The *p*-value of each cuboid block of 2D velocity data was calculated by one-sample *t*-test using those pooled velocity vectors’ projected lengths onto the direction of their averaged vector. The null hypothesis is that there is no cytoplasmic flow and particles merely exhibit Brownian motions. The usage of the color map helps make those significant flow vectors more prominent for better display. Finally, we drew an imaginary embryo cell contour to represent the averaged cell boundary, and removed those data points outside the drawn boundary.

To compare the computational fluid mechanics simulations to the experimental data, we performed analogous projections and averaging. For the simulations, we calculated projected 2D vector fields of fluid flow on 2-μm by 2-μm grids by averaging the continuous velocity field in overlapping domains. For simulations from prometaphase to late anaphase (Figure 3B and 3C, Figure 4E and 4F, and Figure 4H and 4I), we used a 4-μm(x) by 4-μm(y) by 6-μm(z) cuboid averaging domain. For simulations at the PNC stage (Figure 5C and 5D), we used a 4 μm(x) by 4 μm(y) by 8 μm(z) cuboid averaging domain.

### • DATA AND CODE AVAILABILITY

Further information and requests for data and codes should be directed to the Lead Contact, Hai-Yin Wu.

## SUPPLEMENTAL INFORMATION

**Figure S1.**
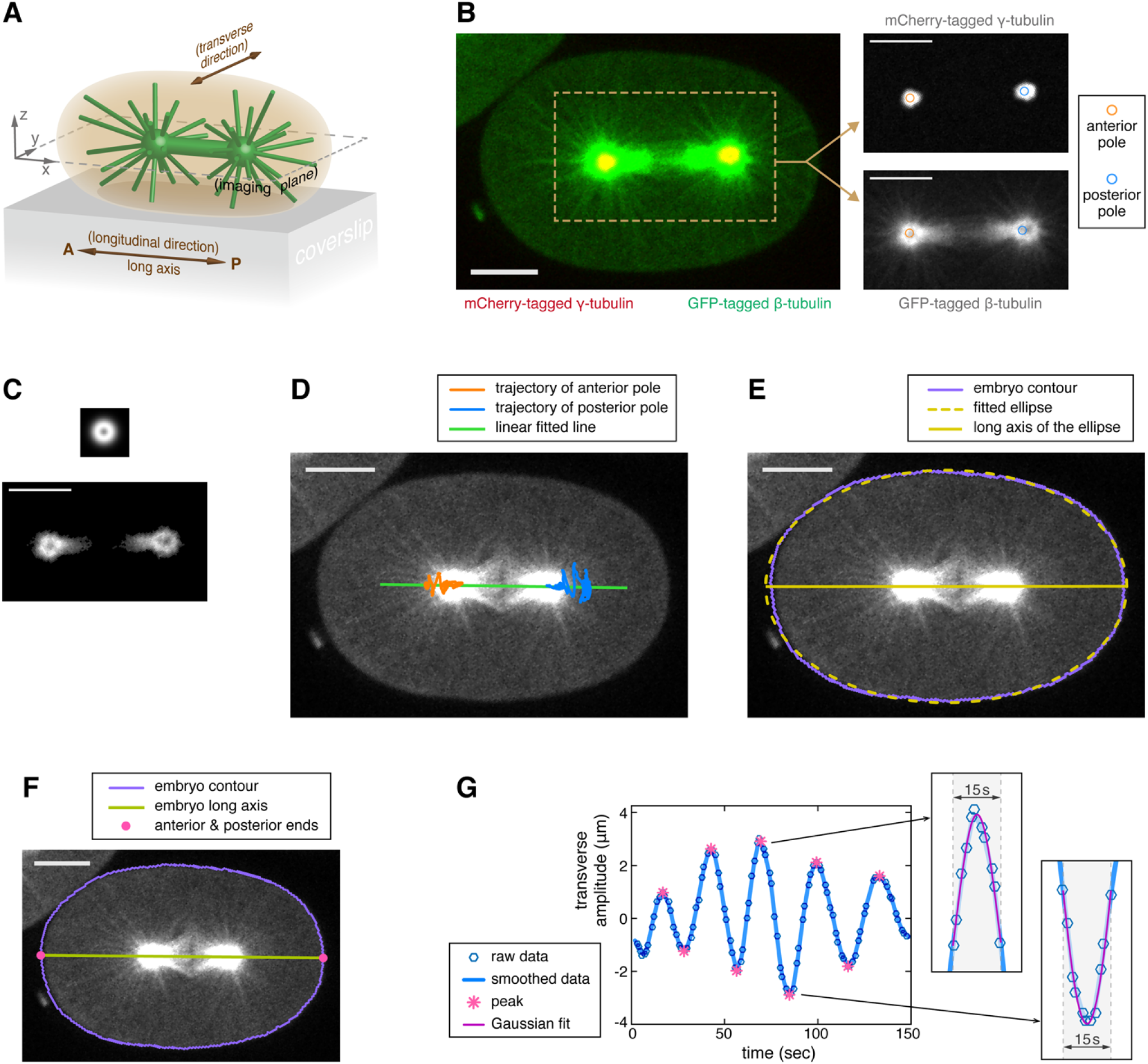
Orientation Terminology, Image Analysis and Centrosome Motion Quantification. (A) The configuration and the terminology of embryo orientation used in this study. The mitotic spindle is simply illustrated by the green dumbbell, while astral microtubules (MTs) by slender green sticks. The confocal imaging plane, which is parallel to the coverslip, is defined as the x-y plane, with the z = 0 plane falling at the center of the spindle or the pronuclear complex (PNC). The embryo’s anterior(A)-to-posterior(P) direction is assigned as the longitudinal direction (x-axis), with the embryo’s posterior end facing the positive x-direction. (B and C) An example of the centrosome-tracking image. (B) The 2-color merged image of an *C. elegans* embryo expressing mCherry-tagged *γ*-tubulin (red) and GFP-tagged β-tubulin (green). The two separate images of the dashed-line region are shown in the right panel. The upper one is *γ*-tubulin-labeling image with its labeled centrosome locations derived by MATLAB Particle Tracking Code Repository by Daniel Blair and Eric Dufresne. The lower one is β-tubulin-labeling image, while its centrosome information is derived from the particle tracking code plus the second-step correlation method. (C) (top) The square image patch with the artificial ring structure mimicking a centrosome. (bottom) An example of the post-processed (background-subtraction and thresholding) β-tubulin-labeling image used for correlation calculation (D and E) The information of the embryo long axis (or longitudinal direction) can be extracted automatically in 2 ways: (D) through linear fitting on the trajectory data of the anterior and posterior centrosomes, or (E) through fitting the automatic detected embryo contour with a simple ellipse. (F) An example of automatic detected embryo contour and embryo anterior/posterior ends. (G) An example of oscillation peak detection. The crests/troughs (or the peak points) of the transverse oscillation were calculated by (1) first finding the local maxima/minima from the smoothed amplitude data, and then (2) fitting the raw amplitude data, which fall within the 15-second windows around the detected maxima/minima, by Gaussian function (see the zoom-in blocks). The derived peak points are indicated by pink asterisks. All scale bars in images: 10 μm.

**Figure S2.**
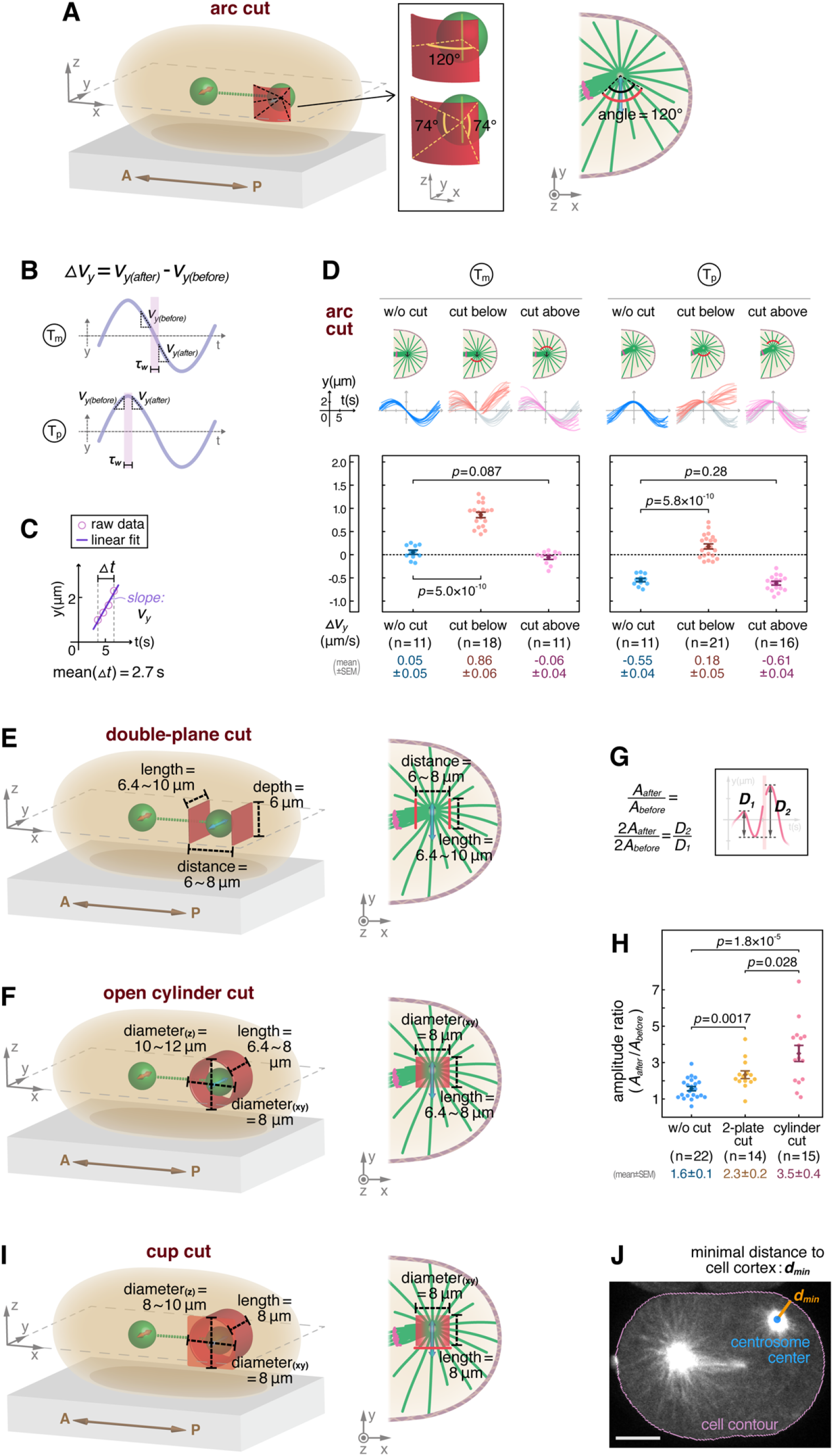
Dimensions of Laser Ablations Performed during Transverse Spindle Oscillations and the Results Quantifications. In 3D schematics of ablation dimensions (A, E, F, and I), centrosomes are represented by two green balls and the spindle body is simplified into the green dashed line. The anterior(A)-to-posterior(P) directions (embryo polarity) are indicated by brown arrows. The orange and blue arrows displayed on centrosomes imply the transverse oscillations of the anterior and posterior centrosomes respectively. The posterior part of the corresponding x-y imaging mid-plane (2D view from the top) is exhibited on the right of each 3D schematic, with the spindle and astral MTs sketched in green and chromosomes in magenta. All ablation geometrics are portrayed in red for both 2D and 3D schematics. (A) Dimensions of the arc cuts in Figure 1E, 1F, 1H, and 1J (all performed with the same angular span stated here). (B and C) Calculation of centrosome velocity change (*Δv_y_*) for arc cuts. (B) Definition of *Δv_y_* at timing T_m_ and T_p_ respectively. And *τ_w_* is the sandwiched window (for ablated embryos: this window is the ablation execution time) between the two intervals *Δt* used for calculating centrosome transverse velocity *v_y(before)_* and *v_y(after)_*. (C) The raw data of centrosome position falling within the interval *Δt* are used to calculate transverse (y-)velocity by linear fitting. (D) The *Δv_y_* data for control and arc-cut embryos. (left) The data for Figure 1E-1F (T_m_), and (right) the data for Figure 1H-1I (T_p_). (E) A schematic of the double-(y-z-)plane cut and their detailed dimensions. (F) Dimensions of the open cylinder cuts in Figure 2A-2E. (G) Measurement on oscillation amplitude change by calculating the amplitude ratio *A_after_/A_before_* (mentioned in Figure 2E). The notation *A* means oscillation transverse amplitude, and *D* stands for peak-to-valley transverse amplitude/displacement. The oscillation example (pink curve) is the one from Figure 2D (open cylinder cut). The ablation execution is highlighted by the light pink vertical strip. (H) The comparison of the amplitude ratios among uncut, double-plane-cut, and open-cylinder-cut embryos. (I) Dimensions of the cup cuts in Figure 2F-2I. (J) An example image of calculating the minimum distance to the cell cortex (mentioned in Figure 2I). The minimum distance, from the posterior centrosome to the cell cortex, of each embryo, is extracted from the time frame with minimal *d_min_*. (Scale bar: 10 μm) All error bars are plotted using standard error of the mean (mean ± SEM). And all displayed *p*-values are calculated by two-tailed Student’s *t*-test.

**Figure S3.**
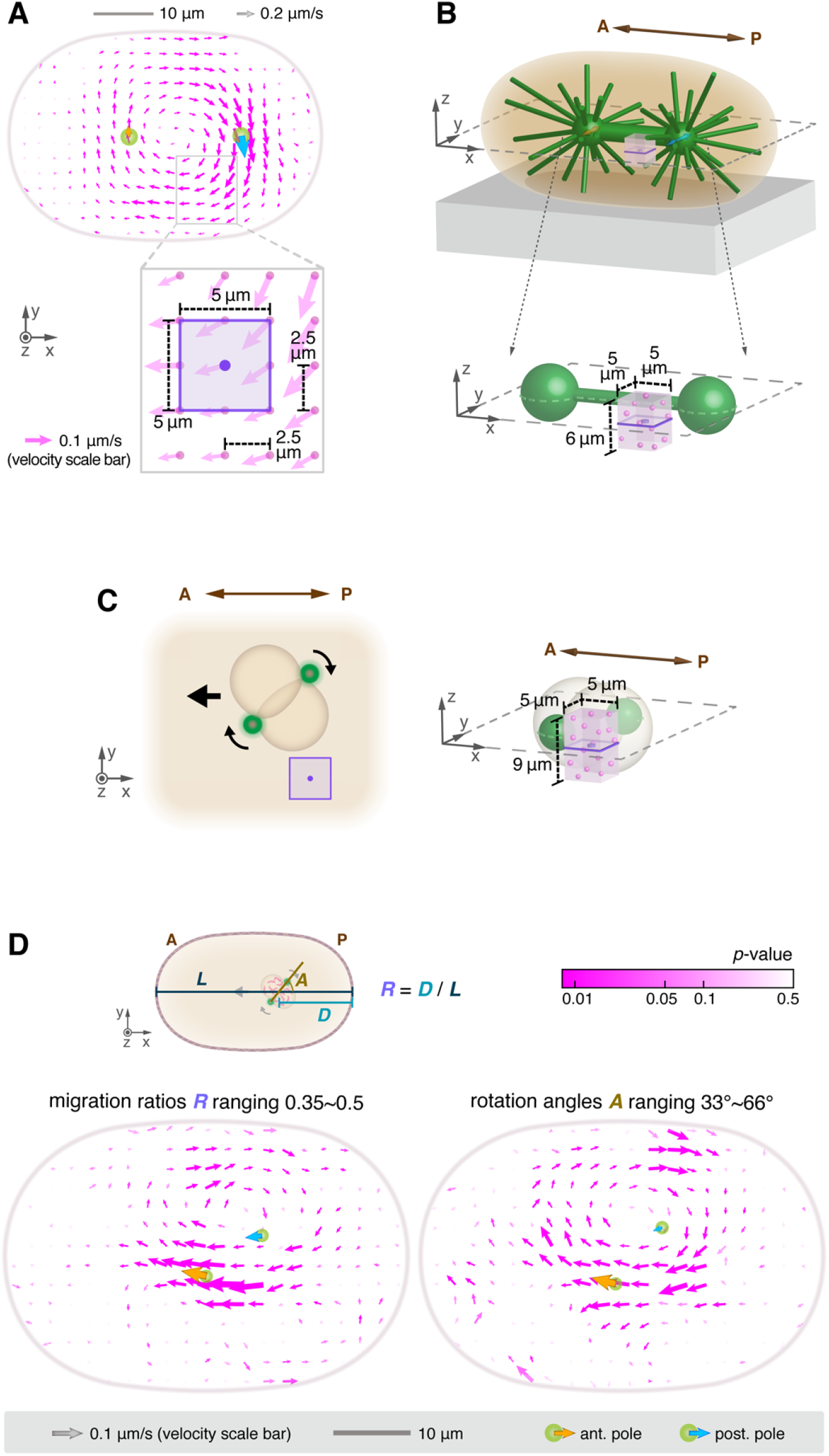
Averaging Operations for Experimental Cytoplasmic Fluid Flow. (A) The fluid flow vector field is displayed by 2D vectors (magenta arrows) plotted on grid points with 2.5-μm x-spacing and 2.5-μm y-spacing (the bottom zoom-in panel). After proper alignment and data grouping, each velocity vector represents the mean velocity of the fluorescent nanodiamonds (FNDs) located within the center grid point’s surrounding averaging domain (the purple square surrounding the purple dot). (B and C) Schematics of the averaging cuboid domain (purple transparent cuboid) with the grid point located at the center of the cuboid. Centrosomes are illustrated by green balls, and the magenta particles inside cuboids represent tracked FNDs. The anterior(A)-to-posterior(P) directions are indicated by brown arrows. (B) For prometaphase to late anaphase, the averaging domain is a 5-μm(x) by 5-μm(y) by 6-μm(z) cuboid. Spindles are aligned onto the middle x-y plane (z = 0), at which plane the mean 2D vector field is plotted. And the purple solid frame (3D view) is the purple square in Figure S3A. (C) An exception for PNC stage, the averaging cuboid measures 5 μm(x) by 5 μm(y) by 9 μm(z) (larger size in the z-direction). The purple square in the left panel is the purple solid frame (3D view) in the right panel. (D) Definition of pronuclear migration/centration and rotation progression. (left) The averaged pronuclear fluid data with migration ratio *R* = 0.35 ~ 0.5 (the same one as Figure 5E), and (right) the data with rotation angle *A* = 33° ~ 66°.

**Figure S4.**
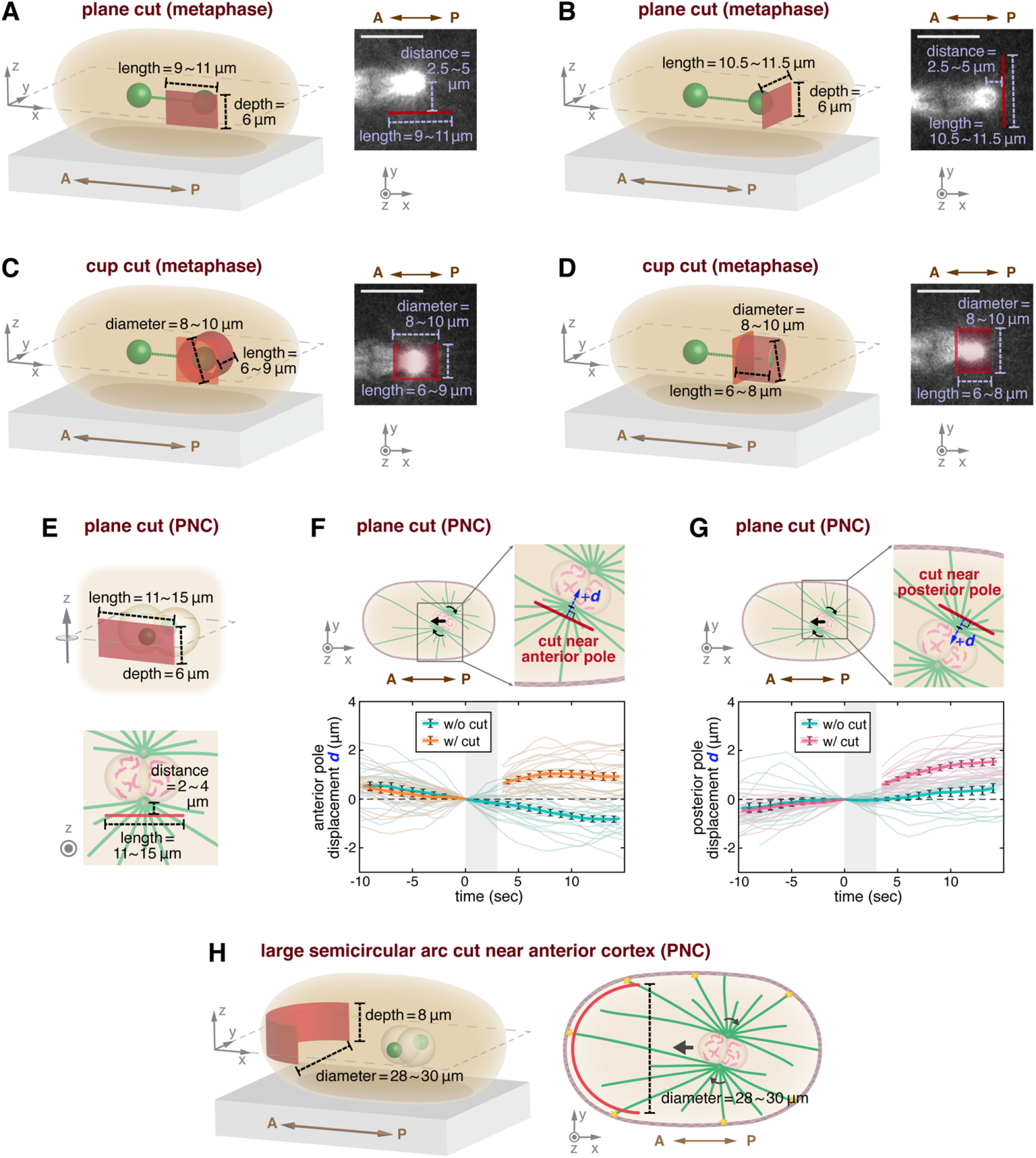
Dimensions of Laser Ablation Experiments Performed in Metaphase and Pronuclear Complex (PNC) Stage. Centrosomes are presented by green balls in all 3D schematics. Centrosomes and astral MTs are sketched in green and chromosomes in magenta in all 2D schematics. All ablation geometrics are portrayed in red. The anterior(A)-to-posterior(P) directions are indicated by brown arrows. (A and B) Metaphase: dimensions of the plane cuts in Figure 4A and 4B respectively. (C and D) Metaphase: dimensions of the cup cuts in Figure 4C and 4D respectively. (E) PNC stage: dimensions of the plane cuts in Figure 5A and 5B. (F and G) PNC stage: (F) the plane cuts near the anterior or leading centrosomes, and (G) the plane cuts near the posterior or trailing centrosomes. Both upper parts are the schematics showing the definition of centrosome center displacement *d*, which is the relative orthogonal (away from the plane cuts) displacement from the locus right before cutting. The black arrows indicate the progression of pronuclear migration and rotation. The mean displacement results with SEM error bars, including uncut and cut embryos, are shown in both lower parts. Their raw data are plotted semi-transparently in the back layer. And the data in main Figure 5B is the combination of these two data sets (F and G), including both plane cuts near the leading/trailing centrosomes. (H) PNC stage: dimensions of the large semicircular arc cuts (near anterior cortices) in Figure 5F-5H. All scale bars in images: 10 μm.

**Figure S5.**
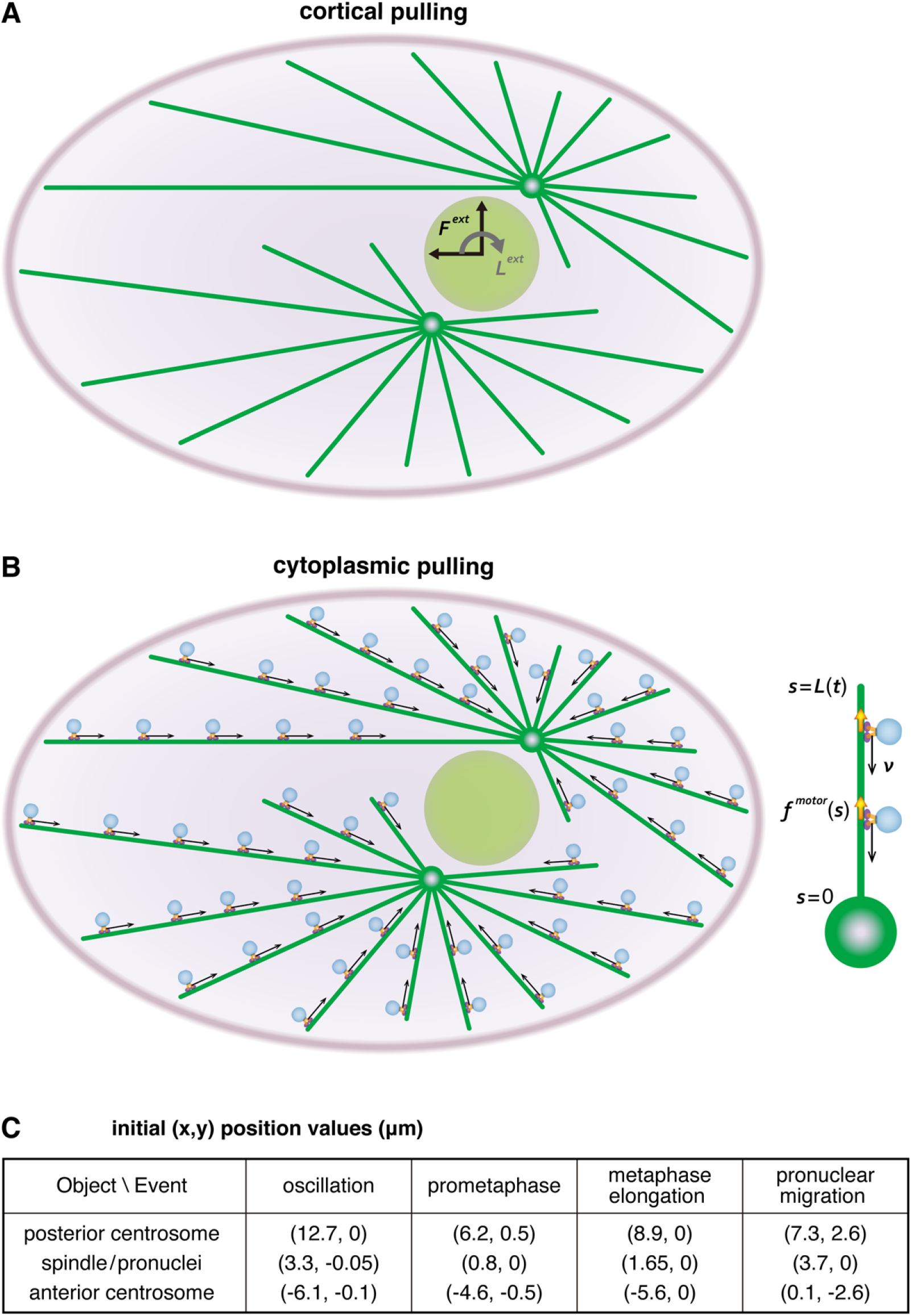
Models of the Force Transduction Mechanisms and Initial Position Values in Computational Fluid Dynamics Simulations. (A and B) Schematics for the computational models of the force transduction mechanisms. The figure shows the PNC stage (see Figure 5E for the corresponding experiment). Our simulations have five structural elements: the cell cortex (gray purple ellipses), the spindle/pronuclei (light green circles), the centrosomes (green circles), elastic MTs (green lines) and the cytoplasm, the fluid filling inside the cortex. (A) *Cortical pulling*. Due to the cortical pulling forces, MTs remain straight and the forces directly act on the PNC without any loss for the MT bending. Hence, in a short time horizon, the cytoplasmic flow arises from the translation and rotation of the pronuclei-centrosome-MT complex. We model that mechanism by applying an external force ***F**^ext^* and torque ***L**^ext^* on the PNC. (B) *Cytoplasmic pulling*. Cytoplasmic dynein motors attach to the MTs and walk towards the centrosomes with velocity ***v*** (black arrow). As they do so, they apply a pulling force on the MTs in the opposite direction to their motion. We model the force applied by the motors by a continuum model: ***f**^motor^*(*s*). (C) The initial (x,y) positions (in μm) of the spindle’s (or pronuclei’s) and the centrosomes’ centers in the simulations. All objects are at z = 0. We decided on those values based on the experiments (see Figure 3E for oscillation, Figure 4G for prometaphase, Figure 4J for metaphase elongation, and Figure 5E for the pronuclear migration).

**Figure S6.**
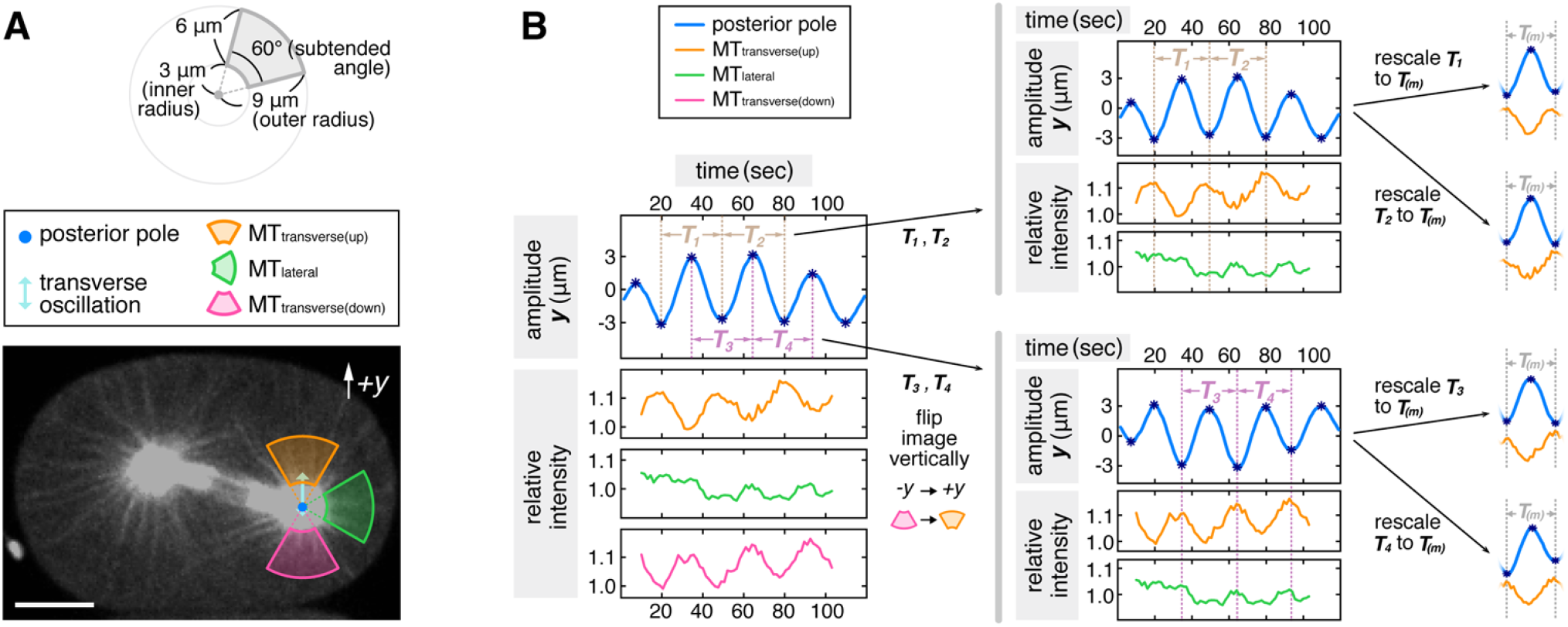
Quantification of Astral Microtubules (MTs) Intensity during Transverse Oscillations. (A) One snapshot from the example GFP::β-tubulin time-series with the assigned transverse and lateral annular-sector regions shown (scale bar: 10 μm). The dimensions of the annular sectors are depicted on the top. (B) Detailed information about data alignment for intensity measurement. Please refer to STAR Methods for detailed operations.

**Table S1.**
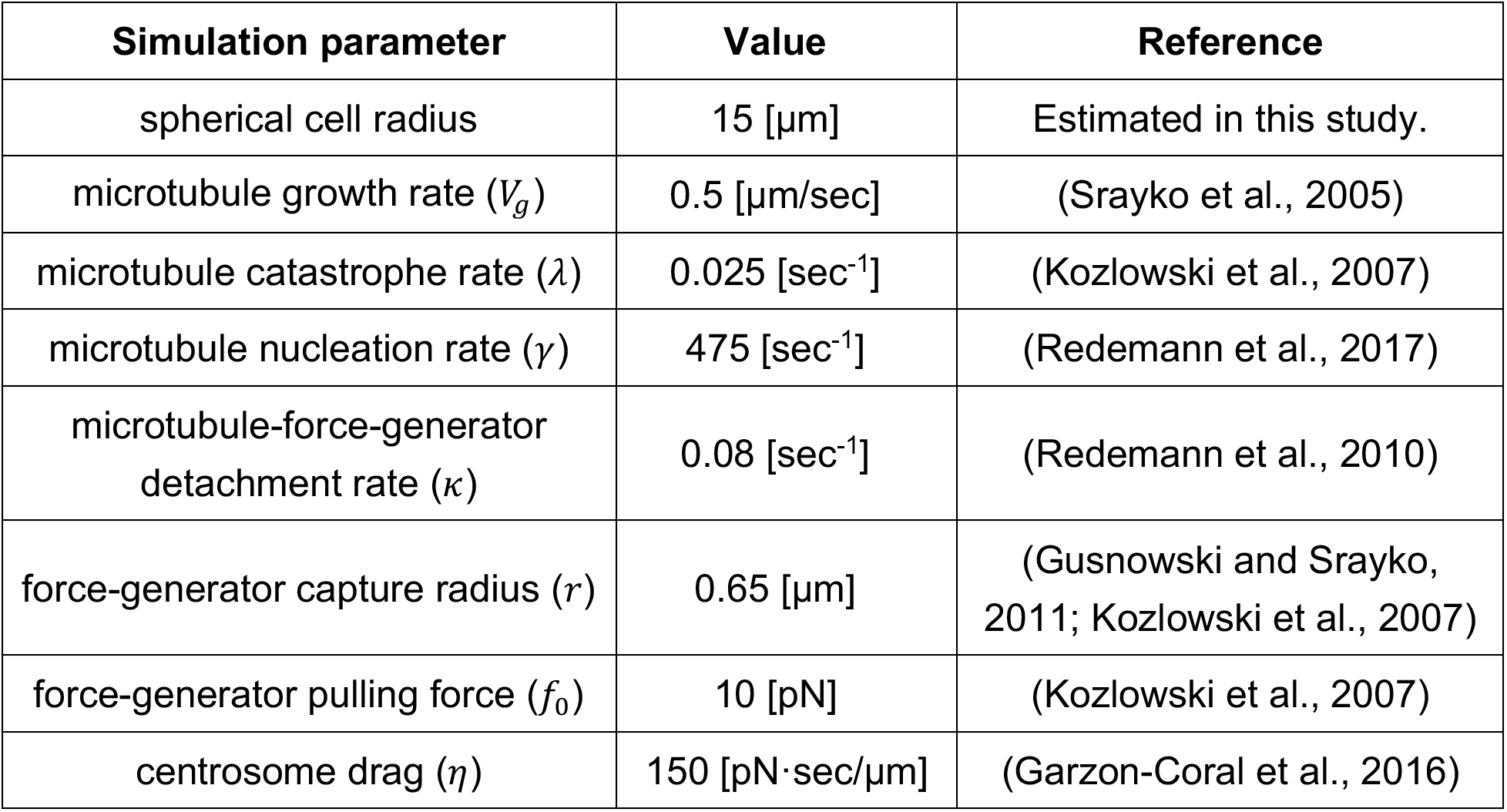
Parameters for the Cortical Pulling Coarse-grained Model, Related to Figure 4.

**Movie S1. Centrosome-positioning Events in an *Caenorhabditis elegans* Early Embryo and the Transverse Spindle Oscillation**

**Movie S2. Arc cuts on Transverse Astral Microtubules (MTs) of Posterior Centrosomes at Two Different Times of Spindle Oscillations**

**Movie S3. Open Cylinder Cuts and Cup Cuts around Posterior Centrosomes During Spindle Oscillations**

**Movie S4. Microinjected Passivated Fluorescent Nanodiamonds (FNDs) in a *Caenorhabditis elegans* Early Embryo**

**Movie S5. Plane Cuts and Cup Cuts around Posterior Centrosomes in Metaphase**

**Movie S6. Plane Cuts around Centrosomes and large Semicircular Arc Cuts Close to Anterior Cortices in Pronuclear Migration and Rotation**

**Movie S7. Measurement of Astral Microtubules (MTs) Intensity within the Annular-sector Regions during Spindle Oscillations**

## ACKNOWLEDGEMENTS

Worm strains were provided by Marie Delattre and the Caenorhabditis Genetics Center. M.J.S. acknowledges support by the National Science Foundation under awards DMR-1420073 (NYU MRSEC), and DMR-2004469. D.J.N. acknowledges the NSF grant DBI-1919834 and the NIH grant 1R01GM104976-01. We are grateful to Che-Hang Yu who assisted with the optical setup; Chia-Yi Fang for advice on FND surface treatment. We also like to thank Reza Farhadifar for helpful discussion over the whole project and advice on modeling; Yuan-Nan Young for assistance in a technical calculation; Joe Howard for helpful discussions.

## AUTHOR CONTRIBUTIONS

H.W., D.J.N., and M.J.S. designed research. H.W. developed the FESLA setup. H.W. conducted all experiments and data analysis. H.C. provided support for fluorescent nanodiamond experiments. G.K., E.N., and M.J.S. worked on large-scale fluid mechanics simulations. M.J.S and D.J.N. derived and analyzed the coarse-grained model. H.W., D.J.N., M.J.S., E.N, and G.K. wrote the manuscript.

## REFERENCES

Bringmann, H., Cowan, C. R., Kong, J. & Hyman, A. A. 2007. LET-99, GOA-1/GPA-16, and GPR-1/2 are required for aster-positioned cytokinesis. Curr Biol, 17, 185–91.

Chang, Y.-R., Lee, H.-Y., Chen, K., Chang, C.-C., Tsai, D.-S., Fu, C.-C., Lim, T.-S., Tzeng, Y.-K., Fang, C.-Y. & Han, C.-C. 2008. Mass production and dynamic imaging of fluorescent nanodiamonds. Nature nanotechnology, 3, 284–288.

Chung, S. H., Clark, D. A., Gabel, C. V., Mazur, E. & Samuel, A. D. 2006. The role of the AFD neuron in C. elegans thermotaxis analyzed using femtosecond laser ablation. BMC neuroscience, 7, 30.

Chung, S. H. & Mazur, E. 2009. Surgical applications of femtosecond lasers. Journal of biophotonics, 2, 557–572.

Coffman, V. C., Mcdermott, M. B., Shtylla, B. & Dawes, A. T. 2016. Stronger net posterior cortical forces and asymmetric microtubule arrays produce simultaneous centration and rotation of the pronuclear complex in the early Caenorhabditis elegans embryo. Molecular biology of the cell, 27, 3550–3562.

Colombo, K., Grill, S. W., Kimple, R. J., Willard, F. S., Siderovski, D. P. & Gonczy, P. 2003. Translation of polarity cues into asymmetric spindle positioning in Caenorhabditis elegans embryos. Science, 300, 1957–61.

Cowan, C. R. & Hyman, A. A. 2004. Asymmetric cell division in C. elegans: cortical polarity and spindle positioning. Annu. Rev. Cell Dev. Biol., 20, 427–453.

Crocker, J. C. & Grier, D. G. 1996. Methods of digital video microscopy for colloidal studies. Journal of colloid and interface science, 179, 298–310.

Daniels, B. R., Masi, B. C. & Wirtz, D. 2006. Probing single-cell micromechanics in vivo: the microrheology of C. elegans developing embryos. Biophys J, 90, 4712–9.

Dogterom, M. & Leibler, S. 1993. Physical aspects of the growth and regulation of microtubule structures. Physical review letters, 70, 1347.

Du, Q. & Macara, I. G. 2004. Mammalian Pins is a conformational switch that links NuMA to heterotrimeric G proteins. Cell, 119, 503–16.

Farhadifar, R., Yu, C.-H., Fabig, G., Wu, H.-Y., Stein, D. B., Rockman, M., Müller-Reichert, T., Shelley, M. J. & Needleman, D. J. 2020. Stoichiometric interactions explain spindle dynamics and scaling across 100 million years of nematode evolution. Elife, 9, e55877.

Foe, V. E. & Von Dassow, G. 2008. Stable and dynamic microtubules coordinately shape the myosin activation zone during cytokinetic furrow formation. J Cell Biol, 183, 457–70.

Fu, C.-C., Lee, H.-Y., Chen, K., Lim, T.-S., Wu, H.-Y., Lin, P.-K., Wei, P.-K., Tsao, P.-H., Chang, H.-C. & Fann, W. 2007. Characterization and application of single fluorescent nanodiamonds as cellular biomarkers. Proceedings of the National Academy of Sciences, 104, 727–732.

Gabel, C. V., Antoine, F., Chuang, C.-F., Samuel, A. D. & Chang, C. 2008. Distinct cellular and molecular mechanisms mediate initial axon development and adult-stage axon regeneration in C. elegans. Development, 135, 1129–1136.

Gabel, C. V., Gabel, H., Pavlichin, D., Kao, A., Clark, D. A. & Samuel, A. D. 2007. Neural circuits mediate electrosensory behavior in Caenorhabditis elegans. Journal of Neuroscience, 27, 7586–7596.

Galli, M., Munoz, J., Portegijs, V., Boxem, M., Grill, S. W., Heck, A. J. & Van Den Heuvel, S. 2011. aPKC phosphorylates NuMA-related LIN-5 to position the mitotic spindle during asymmetric division. Nat Cell Biol, 13, 1132–8.

Gao, Y. & Kilfoi, M. L. 2009. Accurate detection and complete tracking of large populations of features in three dimensions. Optics express, 17, 4685–4704.

Garzon-Coral, C., Fantana, H. A. & Howard, J. 2016. A force-generating machinery maintains the spindle at the cell center during mitosis. Science, 352, 1124–7.

Gotta, M., Dong, Y., Peterson, Y. K., Lanier, S. M. & Ahringer, J. 2003. Asymmetrically Distributed C. elegans Homologs of AGS3/PINS Control Spindle Position in the Early Embryo. Current Biology, 13, 1029–1037.

Götz, T. 2001. Interactions of fibers and flow: asymptotics, theory and numerics, dissertation. de.

Goulding, M. B., Canman, J. C., Senning, E. N., Marcus, A. H. & Bowerman, B. 2007. Control of nuclear centration in the C. elegans zygote by receptor-independent Galpha signaling and myosin II. J Cell Biol, 178, 1177–91.

Grill, S. W., Gonczy, P., Stelzer, E. H. & Hyman, A. A. 2001. Polarity controls forces governing asymmetric spindle positioning in the Caenorhabditis elegans embryo. Nature, 409, 630–3.

Grill, S. W., Howard, J., Schaffer, E., Stelzer, E. H. & Hyman, A. A. 2003. The distribution of active force generators controls mitotic spindle position. Science, 301, 518–21.

Grill, S. W., Kruse, K. & Jülicher, F. 2005. Theory of mitotic spindle oscillations. Physical review letters, 94, 108104.

Gusnowski, E. M. & Srayko, M. 2011. Visualization of dynein-dependent microtubule gliding at the cell cortex: implications for spindle positioning. J Cell Biol, 194, 377–86.

Johnson, R. E. 1980. An improved slender-body theory for Stokes flow. Journal of Fluid Mechanics, 99, 411–431.

Keller, J. B. & Rubinow, S. I. 1976. Slender-body theory for slow viscous flow. Journal of Fluid Mechanics, 75, 705–714.

Kimura, A. & Onami, S. 2005. Computer simulations and image processing reveal length-dependent pulling force as the primary mechanism for C-elegans male pronuclear migration. Developmental Cell, 8, 765–775.

Kimura, K. & Kimura, A. 2011. Intracellular organelles mediate cytoplasmic pulling force for centrosome centration in the Caenorhabditis elegans early embryo. Proc Natl Acad Sci U S A, 108, 137–42.

Kiyomitsu, T. & Cheeseman, I. M. 2012. Chromosome- and spindle-pole-derived signals generate an intrinsic code for spindle position and orientation. Nat Cell Biol, 14, 311–7.

Knoblich, J. A. 2010. Asymmetric cell division: recent developments and their implications for tumour biology. Nat Rev Mol Cell Biol, 11, 849–60.

Kotak, S., Busso, C. & Gönczy, P. 2012. Cortical dynein is critical for proper spindle positioning in human cells. J Cell Biol, 199, 97–110.

Kozlowski, C., Srayko, M. & Nedelec, F. 2007. Cortical microtubule contacts position the spindle in C. elegans embryos. Cell, 129, 499–510.

Krueger, L. E., Wu, J. C., Tsou, M. F. & Rose, L. S. 2010. LET-99 inhibits lateral posterior pulling forces during asymmetric spindle elongation in C. elegans embryos. J Cell Biol, 189, 481–95.

Laan, L., Pavin, N., Husson, J., Romet-Lemonne, G., Van Duijn, M., López, M. P., Vale, R. D., Jülicher, F., Reck-Peterson, S. L. & Dogterom, M. 2012. Cortical dynein controls microtubule dynamics to generate pulling forces that position microtubule asters. Cell, 148, 502–514.

Longo, F. J. & Anderson, E. 1968. The fine structure of pronuclear development and fusion in the sea urchin, Arbacia punctulata. J Cell Biol, 39, 339–68.

Malhotra, D. & Biros, G. 2015. PVFMM: A parallel kernel independent FMM for particle and volume potentials. Communications in Computational Physics, 18, 808–830.

Meaders, J. L. & Burgess, D. R. 2020. Microtubule-Based Mechanisms of Pronuclear Positioning. Cells, 9.

Minc, N., Burgess, D. & Chang, F. 2011. Influence of cell geometry on division-plane positioning. Cell, 144, 414–26.

Mochalin, V. N., Shenderova, O., Ho, D. & Gogotsi, Y. 2012. The properties and applications of nanodiamonds. Nature nanotechnology, 7, 11.

Mohan, N., Chen, C.-S., Hsieh, H.-H., Wu, Y.-C. & Chang, H.-C. 2010. In vivo imaging and toxicity assessments of fluorescent nanodiamonds in Caenorhabditis elegans. Nano letters, 10, 3692–3699.

Nazockdast, E., Rahimian, A., Needleman, D. & Shelley, M. 2017a. Cytoplasmic flows as signatures for the mechanics of mitotic positioning. Molecular biology of the cell, 28, 3261–3270.

Nazockdast, E., Rahimian, A., Needleman, D. & Shelley, M. 2017b. Cytoplasmic flows as signatures for the mechanics of mitotic positioning. Mol Biol Cell, 28, 3261–3270.

Nazockdast, E., Rahimian, A., Zorin, D. & Shelley, M. 2017c. A fast platform for simulating semi-flexible fiber suspensions applied to cell mechanics. Journal of Computational Physics, 329, 173–209.

Nguyen-Ngoc, T., Afshar, K. & Gonczy, P. 2007. Coupling of cortical dynein and G alpha proteins mediates spindle positioning in Caenorhabditis elegans. Nat Cell Biol, 9, 1294–302.

Pearson, C. G. & Bloom, K. 2004. Dynamic microtubules lead the way for spindle positioning. Nat Rev Mol Cell Biol, 5, 481–92.

Pecreaux, J., Roper, J. C., Kruse, K., Julicher, F., Hyman, A. A., Grill, S. W. & Howard, J. 2006. Spindle oscillations during asymmetric cell division require a threshold number of active cortical force generators. Curr Biol, 16, 2111–22.

Pozrikidis, C. 1992. Boundary integral and singularity methods for linearized viscous flow, Cambridge University Press.

Redemann, S., Baumgart, J., Lindow, N., Shelley, M., Nazockdast, E., Kratz, A., Prohaska, S., Brugués, J., Fürthauer, S. & Müller-Reichert, T. 2017. C. elegans chromosomes connect to centrosomes by anchoring into the spindle network. Nature communications, 8, 1–13.

Reinsch, S. & Gonczy, P. 1998. Mechanisms of nuclear positioning - Commentary. Journal of Cell Science, 111, 2283–2295.

Reinsch, S. & Karsenti, E. 1997. Movement of nuclei along microtubules in Xenopus egg extracts. Curr Biol, 7, 211–4.

Riche, S., Zouak, M., Argoul, F., Arneodo, A., Pecreaux, J. & Delattre, M. 2013. Evolutionary comparisons reveal a positional switch for spindle pole oscillations in Caenorhabditis embryos. Journal of Cell Biology, 201, 653–662.

Roca-Cusachs, P., Conte, V. & Trepat, X. 2017. Quantifying forces in cell biology. Nature cell biology, 19, 742–751.

Rujano, M. A., Sanchez-Pulido, L., Pennetier, C., Le Dez, G. & Basto, R. 2013. The microcephaly protein Asp regulates neuroepithelium morphogenesis by controlling the spatial distribution of myosin II. Nat Cell Biol, 15, 1294–306.

Saad, Y. & Schultz, M. H. 1986. GMRES: A generalized minimal residual algorithm for solving nonsymmetric linear systems. SIAM Journal on scientific and statistical computing, 7, 856–869.

Shinar, T., Mana, M., Piano, F. & Shelley, M. J. 2011. A model of cytoplasmically driven microtubule-based motion in the single-celled Caenorhabditis elegans embryo. Proc Natl Acad Sci U S A, 108, 10508–13.

Siller, K. H. & Doe, C. Q. 2009. Spindle orientation during asymmetric cell division. Nat Cell Biol, 11, 365–74.

Stein, D. B., De Canio, G., Lauga, E., Shelley, M. J. & Goldstein, R. E. 2021. Swirling Instability of the Microtubule Cytoskeleton. Physical Review Letters, 126, 028103.

Su, L.-J., Wu, M.-S., Hui, Y. Y., Chang, B.-M., Pan, L., Hsu, P.-C., Chen, Y.-T., Ho, H.-N., Huang, Y.-H. & Ling, T.-Y. 2017. Fluorescent nanodiamonds enable quantitative tracking of human mesenchymal stem cells in miniature pigs. Scientific reports, 7, 45607.

Tame, M. A., Raaijmakers, J. A., Van Den Broek, B., Lindqvist, A., Jalink, K. & Medema, R. H. 2014. Astral microtubules control redistribution of dynein at the cell cortex to facilitate spindle positioning. Cell Cycle, 13, 1162–70.

Tanimoto, H., Sallé, J., Dodin, L. & Minc, N. 2018. Physical forces determining the persistency and centring precision of microtubule asters. Nature physics, 14, 848–854.

Tornberg, A.-K. & Shelley, M. J. 2004. Simulating the dynamics and interactions of flexible fibers in Stokes flows. Journal of Computational Physics, 196, 8–40.

Tsou, M. F., Hayashi, A., Debella, L. R., Mcgrath, G. & Rose, L. S. 2002. LET-99 determines spindle position and is asymmetrically enriched in response to PAR polarity cues in C. elegans embryos. Development, 129, 4469–81.

Vaijayanthimala, V., Cheng, P.-Y., Yeh, S.-H., Liu, K.-K., Hsiao, C.-H., Chao, J.-I. & Chang, H.-C. 2012. The long-term stability and biocompatibility of fluorescent nanodiamond as an in vivo contrast agent. Biomaterials, 33, 7794–7802.

Valentine, M. T., Perlman, Z. E., Gardel, M. L., Shin, J. H., Matsudaira, P., Mitchison, T. J. & Weitz, D. A. 2004. Colloid surface chemistry critically affects multiple particle tracking measurements of biomaterials. Biophys J, 86, 4004–14.

Vogel, A., Noack, J., Hüttman, G. & Paltauf, G. 2005. Mechanisms of femtosecond laser nanosurgery of cells and tissues. Applied Physics B, 81, 1015–1047.

Von Dassow, G., Verbrugghe, K. J., Miller, A. L., Sider, J. R. & Bement, W. M. 2009. Action at a distance during cytokinesis. J Cell Biol, 187, 831–45.

Walston, T. & Hardin, J. 2010. An agar mount for observation of Caenorhabditis elegans embryos. Cold Spring Harb Protoc, 2010, pdb.prot5540.

Wirtz, D. 2009. Particle-tracking microrheology of living cells: principles and applications. Annu Rev Biophys, 38, 301–26.

Wu, H. Y., Nazockdast, E., Shelley, M. J. & Needleman, D. J. 2017. Forces positioning the mitotic spindle: Theories, and now experiments. Bioessays, 39.

Xie, J. & Minc, N. 2020. Cytoskeleton Force Exertion in Bulk Cytoplasm. Frontiers in Cell and Developmental Biology, 8.

Yu, C.-H., Redemann, S., Wu, H.-Y., Kiewisz, R., Yoo, T. Y., Conway, W., Farhadifar, R., Müller-Reichert, T. & Needleman, D. 2019. Central-spindle microtubules are strongly coupled to chromosomes during both anaphase A and anaphase B. Molecular biology of the cell, 30, 2503–2514.

Zhu, M., Settele, F., Kotak, S., Sanchez-Pulido, L., Ehret, L., Ponting, C. P., Gönczy, P. & Hoffmann, I. 2013. MISP is a novel Plk1 substrate required for proper spindle orientation and mitotic progression. J Cell Biol, 200, 773–87.

